# Generation of kidney ureteric bud and collecting duct organoids that recapitulate kidney branching morphogenesis

**DOI:** 10.1101/2020.04.27.049031

**Authors:** Zipeng Zeng, Biao Huang, Riana K. Parvez, Yidan Li, Jyunhao Chen, Ariel Vonk, Matthew E. Thornton, Tadrushi Patel, Elisabeth A. Rutledge, Albert D. Kim, Jingying Yu, Nuria Pastor-Soler, Kenneth R. Hallows, Brendan H. Grubbs, Jill A. McMahon, Andrew P. McMahon, Zhongwei Li

## Abstract

Kidney organoids model development and diseases of the nephron but not the contiguous epithelial network of the kidney’s collecting duct (CD) system. Here, we report the generation of an expandable, 3D branching ureteric bud (UB) organoid culture model that can be derived from primary UB progenitors from mouse and human fetal kidneys, or generated *de novo* from pluripotent human stem cells. UB organoids differentiate into CD organoids *in vitro*, with differentiated cell types adopting spatial assemblies reflective of the adult kidney collecting system. Aggregating 3D-cultured nephron progenitor cells with UB organoids *in vitro* results in a reiterative process of branching morphogenesis and nephron induction, similar to kidney development. Combining efficient gene editing with the UB organoid model will facilitate an enhanced understanding of development, regeneration and diseases of the mammalian collecting system.

**One sentence summary:** Collecting duct organoids derived from primary mouse and human ureteric bud progenitor cells and human pluripotent stem cells provide an *in vitro* platform for genetic dissection of development, regeneration and diseases of the mammalian collecting system.

## INTRODUCTION

The mammalian kidney contains thousands of nephrons, connected to a highly branched collecting duct (CD) system. Nephrons filter and process the blood to form the primitive urine, which is collected and further refined in the CD system to adjust water, electrolytes and pH and to maintain the homeostasis of tissue fluid^1,2^. The complex and elaborate kidney is largely formed from the reciprocal interactions of two embryonic cell populations: the epithelial ureteric bud (UB); and the surrounding metanephric mesenchyme (MM). Signals from the MM induce the repeated branching of UB, which gives rise to the entire CD network. Meanwhile, signals from the UB induce the MM to form nephrons^3,4^. Given this central role of the UB in kidney organogenesis, defects in UB/CD development often lead to malformation of the kidney, low endowment of nephrons at birth, and congenital anomalies of kidney and urinary tract (CAKUT)^2,4,5^. Thus, a better understanding of kidney branching morphogenesis is needed for *in vitro* efforts towards rebuilding the kidney. It is also required for developing novel preventive, diagnostic and therapeutic approaches for various kidney diseases.

Three-dimensional (3D) multicellular mini-organ structures, or organoids, have broad applications for modeling organ development and disease, and for regenerating organs through cell or tissue replacement therapies^6,7^. Recently, we and others have been able to generate kidney organoids from human pluripotent stem cells^8-11^ or from expandable nephron progenitor cells (NPCs)^12-14^. These organoids have greatly aided studies of the role of nephrons in kidney development and disease^15^. However, we still lack a robust kidney organoid model that can recapitulate UB branching morphogenesis and its maturation into the renal CD network, despite previous efforts relying on primary mouse/rat tissue^16-20^ or human pluripotent stem cells^21-24^.

Here, we report the development of a 3D organoid model that mimics the full spectrum of kidney branching morphogenesis *in vitro*—from the expandable immature UB progenitor stage, to the mature CD stage. These organoids, derived from primary UB progenitor cells and human pluripotent stem cells, are amenable to efficient gene editing, and have broad applications for studying kidney development, regeneration and disease.

## RESULTS

### Expanding mouse UB progenitor cells into 3D branching UB organoids

We previously developed a 3D culture system for the long-term expansion of mouse and human nephron progenitor cells (NPCs), which can generate nephron organoids that recapitulate kidney development and disease^14,25^. UB branching morphogenesis is driven by another kidney progenitor population, the UB progenitor cells (UPCs). We thus set out to establish a culture system for the expansion of UPCs. UPCs are specified around embryonic day 10.5 (E10.5), when the UB starts to invade the MM. UPCs disappear around postnatal day 2 (P2), when nephrogenesis ceases. Self-renewing UPCs reside in the tip region of the branching UB. During their approximately 10-day lifespan, some UPCs migrate out of UB tip niche to the UB trunk, and differentiate into the renal CD network. Other UPCs proliferate and replenish the self-renewing progenitor cell population of the UB tip. *Ret*^*26,27*^ and *Wnt11*^*28*^ have been identified as specific markers for UPCs, and the transgenic reporter mouse strain *Wnt11*-myrTagRFP-IRES-CE (“*Wnt11*-RFP” for short) has been generated to facilitate the real-time tracking of *Wnt11*-expresing cells based on RFP expression, and the lineage tracing of their progeny via a Cre-mediated recombination system^29^.

We employed this *Wnt11*-RFP reporter system as a readout to screen for a culture condition that maintained the progenitor identity of UPCs *in vitro*. T-shaped UBs were manually isolated from E11.5 kidneys of *Wnt11*-RFP mice, and immediately embedded into Matrigel to set up a 3D culture platform that supported epithelial branching. Using this 3D culture format, hundreds of different combinations of growth factors and small molecules were tested, following strategies similar to those we used to establish optimal NPC culture^14^. This screening allowed us to identify a cocktail, which we named “UB culture medium” (UBCM), that maintained self-renewing UPCs as a 3D branching UB organoid (**Fig. 1a**). Under this culture condition, the T-shaped UB formed a rapidly expanding branching epithelial morphology, with uniform *Wnt11*-RFP expression maintained throughout the 3D structure (**Fig. 1b**). Resected *Wnt11*-RFP^+^ UB organoid tips, re-embedded in Matrigel, branched and grew into additional *Wnt11*-RFP^+^ UB organoids. Repetitive passaging and embedding for up to 3 weeks, resulted in over a hundred thousand-fold expansion in the number of cells (**Fig. 1c**). *Wnt11*-RFP levels remained uniform for the first 10 days and progressively dropped thereafter, similar to the normal course of UPCs *in vivo* (**Fig. 1c**). Consistent with the uniform expression of *Wnt11*-RFP in the entire UB organoid, whole-mount immunostaining of cultures after 10 days, confirmed the homogenous expression of UPC markers *Ret, Etv5*, and *Sox9*, as well as broad UB lineage markers *Gata3, Pax2, Pax8, Krt8*, and *Cdh1*, in the UB organoids (**Fig. 1d,e; Extended Data Fig. 1a,b**).

**Figure 1.**
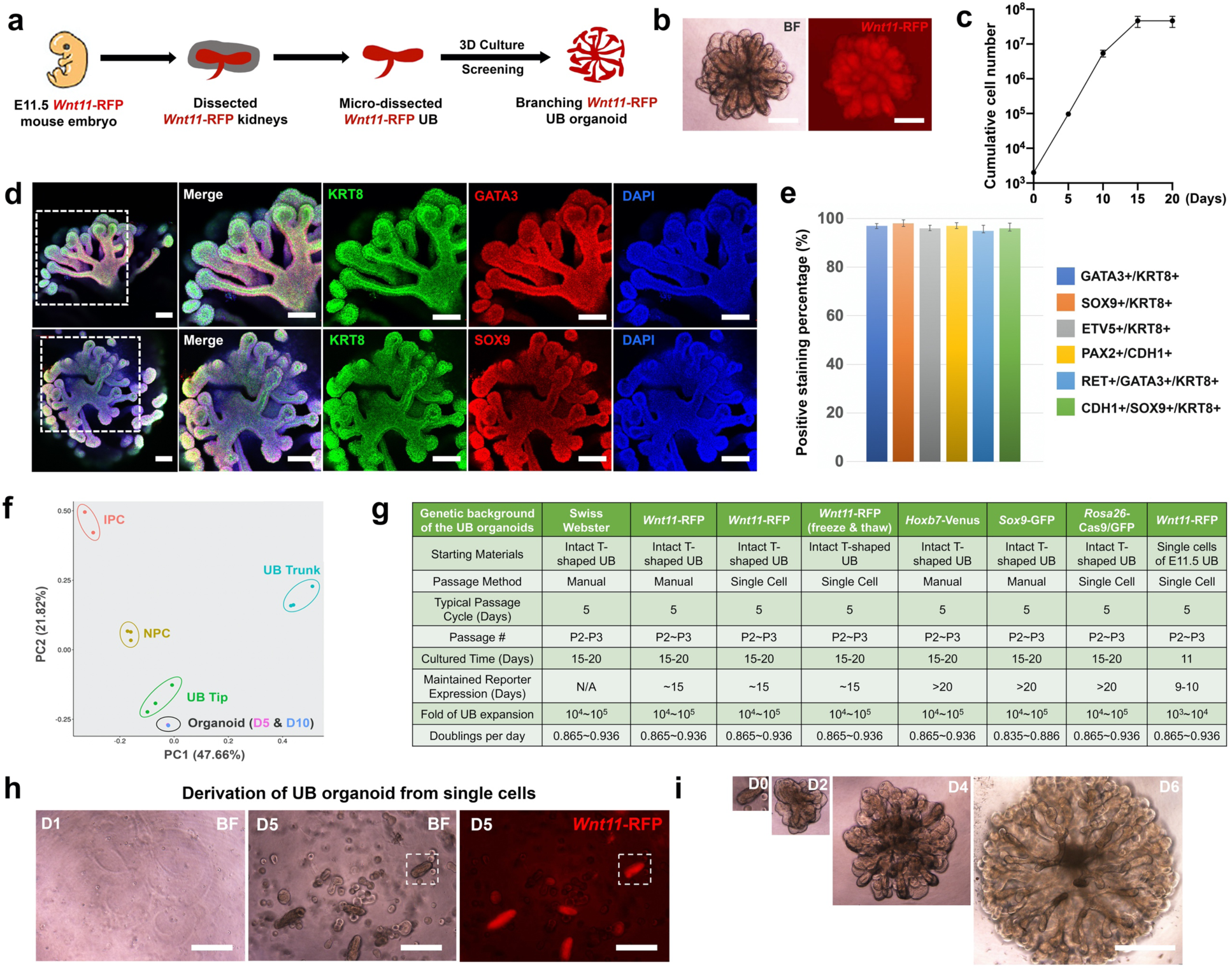
Expansion of mouse UB progenitor cells as 3D branching UB organoids. **a**, Schematic of mouse UB isolation and screening for UB organoid culture condition. **b**, Representative bright field (BF, left panel) and *Wnt11*-RFP (right panel) images of UB organoid. Scale bars, 200 µm. **c**, Cumulative growth curve of UB organoid culture starting from 2000 cells. Data are presented as mean ± s.d. Each time point represents 3 biological replicates. **d**, Whole-mount immunostaining of UB organoid for various UB markers after 10 days of culture. Scale bars, 100 µm. **e**, Quantification of UB organoid immunostaining images. Data are presented as mean ± s.d. **f**, Principal component analysis (PCA) of RNA-seq data. Different colors and oval cycles represent different primary kidney progenitor cell populations or UB organoids cultured for 5 days (D5) and 10 days (D10). Note the large overlap in D5 and D10 organoid samples. **g**, Summary of UB organoid derivation from mouse strains with different genetic backgrounds. **h.** Bright field images showing single UB cells cultured in the UBCM on Day 1 (left panel) and Day 5 (middle panel), as well as *Wnt11-*RFP image on Day 5 (right panel). Scale bars, 200 µm. **i**, Derivation of UB organoid from a single UB cell-derived budding structure as boxed in **h**. Scale bar, 400 µm.

To better define the identity of the UB organoids, we used RNA-seq to profile the transcriptome of the organoids after 5 days and 10 days in culture. These data were compared with prior RNA-seq data for primary UB tip and UB trunk populations^30^, as well as for NPCs and interstitial progenitor cells (IPCs)^31^. Unsupervised clustering (**Extended Data Fig. 1c**) and principle component analysis (PCA) (**Fig. 1f**) placed the cultured UB organoids closer to the primary UB tip samples than to differentiated stalk derivatives of the UB trunk (**Extended Data Fig. 1c**). Taken together, these findings indicate the UB organoid culture system enables a substantial expansion of cells retaining molecular characteristics of UPCs *in vitro*.

Next, we tested whether the UB organoid culture system can be applied to mouse strains other than *Wnt11*-RFP. For this, we successfully derived UB organoids from E11.5 UB from Swiss Webster, a wild-type mouse strain, and from multiple transgenic mouse strains including *Hoxb7*-Venus^32^, *Sox9*-GFP^33^, and *Rosa26*-Cas9/GFP^34^ (**Extended Data Fig. 1d,e** and data not shown). All of these UB organoids retained the typical branching morphology and showed very similar growth rates, compared to *Wnt11*-RFP UB organoids (**Fig. 1g**), indicating the robustness of the 3D/UBCM culture system. Moreover, UB organoids still self-organized into branching organoids after a freeze-thaw process, adding flexibility to the culture system with regards to cell storage (**Extended Data Fig. 1f**).

In determining whether we had developed a synthetic niche for UPC self-renewal, the most stringent test was whether the UBCM culture condition was able to derive UB organoids from single cells. For this purpose, we dissociated E11.5 UBs into single cells before embedding them into Matrigel and culturing in UBCM medium. Around 30% of the single cells self-organized into E11.5 UB-like budding structures within 5 days, though a smaller percentage (3-5%) maintained *Wnt11*-RFP (**Fig. 1h; Extended Data Fig. 1g,h**), an efficiency similar to clonal organoid formation for *Lgr5*^+^ intestinal stem cells^35^. Importantly, the clonally-derived *Wnt11*-RFP^+^ budding structures were identical to intact E11.5 UB-derived organoids in both branching morphology and growth rate (**Fig. 1g-i**). Furthermore, withdrawal of the major medium components from UBCM resulted in either growth arrest (CHIR99021) or rapid loss of *Wnt11*-RFP (all other components), suggesting that each component was essential for optimal UB organoid culture (**Extended Data Fig. 2a-c**). These data, taken together, suggest that UBCM represents a synthetic niche for the *in vitro* expansion of UPCs.

### Screening for conditions to mature UB organoids into CD organoids

The functions of the mature renal CD system are carried out by two major cell populations that are intermingled throughout the entire CD network. The more abundant principal cells (PCs) concentrate the urine and regulate Na^+^/K^+^ homeostasis via water and Na^+^/K^+^ transporters. The less abundant α- and β-intercalated cells (ICs) regulate normal acid-base homeostasis via secretion of H^+^ or HCO3^-^ into the urine. The absence of an *in vitro* system recapitulating PC and IC development in an appropriate 3D context, constrains physiological exploration, disease modeling and drug screening on the renal CD system. With this limitation in mind, we developed a screen to establish conditions supporting the differentiation of CD organoids, assaying expression of *Aqp2*^*36*^ and *Foxi1*^*37*^, definitive markers for PC and IC lineages, respectively, by quantitative reverse transcription PCR (qRT-PCR), following 7 days of culture under variable but defined culture conditions (**Fig. 2a**).

**Figure 2.**
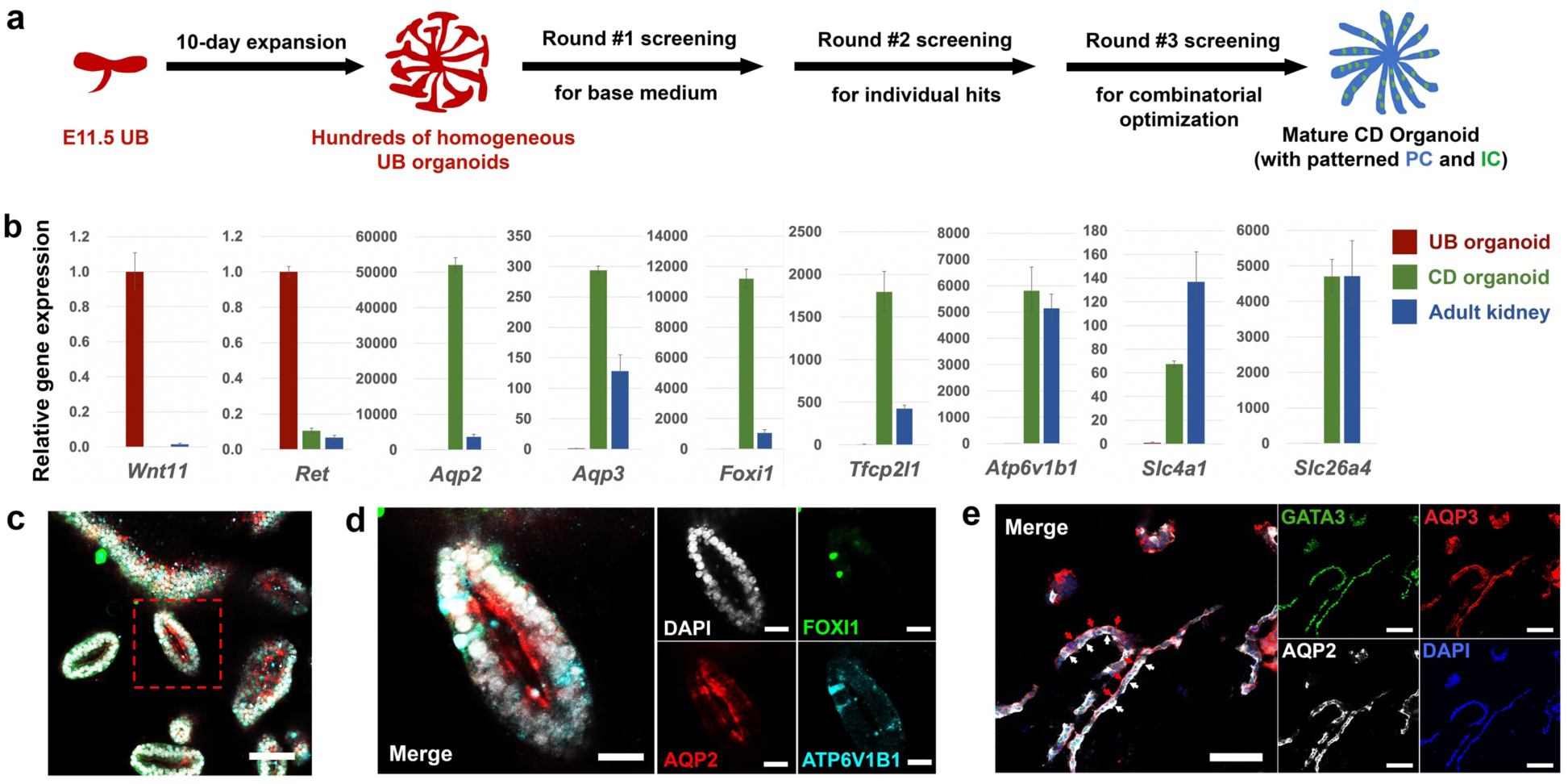
Generation of mature and highly organized CD organoids from UB organoids. **a**, Schematic of the screening strategy for identifying differentiation culture condition to generate CD organoids from UB organoids. **b**, qRT-PCR analyses of UB (red) and CD (green) organoids for UB progenitor markers *Wnt11* and *Ret*, PC markers *Aqp2* and *Aqp3*, IC markers *Foxi1, Atp6v1b1, Slc4a1*, and *Slc26a4*, and *Tfcp2l1* that is expressed in both PC and IC^42^. Adult mouse kidney (blue) was used as control. Data are presented as mean ± s.d. Each column represents 3 technical replicates. **c-e**, Whole-mount (**c, d**) and cryosection (**e**) immunostaining analyses of the CD organoids for broad ureteric lineage marker GATA3, PC markers AQP2 and AQP3, and IC markers FOXI1 and ATP6V1B1. (**d**) shows the higher-power images for the boxed region in (**c**). In (**e**), white arrows indicate AQP2 expression lining the apical plasma membrane (facing the organoid lumen) of the epithelia, while red arrows indicate AQP3 expression lining the basolateral membrane. Scale bars, **c**, 100 µm; **d**, 25 µm; **e**, 50 µm.

In a 1^st^ round of screening, we determined the base condition in which minimal growth factors/small molecules sustained the survival of the organoids and permitted their differentiation. The base medium used for UBCM – hBI^14^ – was tested, together with the commercially available APEL medium for sustaining kidney organoid generation^9^. Combinations of FGF9, EGF and Y27632 (empirically determined) were tested, together with the two different base media (**Extended Data Fig. 3a**). After 7 days of differentiation in the various conditions, we observed that the hBI+FGF9+Y27632 condition enabled the survival of the organoid and permitted spontaneous basal differentiation, as shown by the modest elevation of both *Aqp2* and *Foxi1* (**Extended Data Fig. 3b,c**).

To enhance the efficiency of differentiation, we carried out a 2^nd^ round of screening identifying molecules that strongly induced the expression of *Aqp2* and/or *Foxi1* under the hBI+FGF9+Y27632 condition. Agonists or antagonists targeting major developmental pathways (TGF-β, BMP, Wnt, FGF, Hedgehog and Notch) were tested, together with hormonal inputs known to regulate PC or IC activity (aldosterone and vasopressin). BMP7, DAPT (a Notch pathway inhibitor), JAKI (JAK inhibitor I) and PD0325901 (MEK inhibitor) dramatically increased both *Aqp2* and *Foxi1* expression, while JAG-1^38^ (Notch agonist) and aldosterone led to a preferential increase in *Foxi1* expression, and vasopressin to enhanced *Aqp2* expression (**Extended Data Fig. 3d-f**). In a 3^rd^ round of screening, testing of various combinations of these factors led to the identification of an optimized CD differentiation medium (CDDM) supplemented with FGF9, Y27632, DAPT, PD0325901, aldosterone and vasopressin.

### Generating mature and highly organized CD organoids from UB organoids

Seven days of UB organoid culture in CDDM resulted in a marked decrease in expression of the UPC genes (*Wnt11* and *Ret*) and a concomitant elevation in the expression of PC-specific water transporter encoding genes (*Aqp2* and *Aqp3*^36^) and IC-specific transcription factor (*Foxi1*), proton pump (*Atp6v1b1*^39^) and Cl^-^/HCO3^-^ exchangers (*Slc4a1*/*Ae1*^40^, α-IC-specific; *Slc26a4*/*Pendrin*^41^ β-IC specific) (**Fig. 2b**). Immunostaining confirmed the presence of AQP2, AQP3, FOXI1 and TFCP2L1^42^ in CD organoids (**Fig. 2c-e; Extended Data Fig. 3g,h**). Differentiating CD organoids displayed a clear lumen (**Fig. 2c**), and the organization of PC and IC cell types reflected that of the kidney CD: AQP2^+^/AQP3^+^ PCs comprised 70-90% of the cells, and FOXI1^+^/ATP6V1B1^+^/TFCP2L1^+^/KIT^+43,44^ ICs were widely dispersed, solitary cells (**Fig. 2c,d; Extended Data Fig. 3g,h**). Further, AQP2 and AQP3 showed a differential subcellular localization, AQP2 to the apical luminal facing surface and AQP3 to basolateral plasma membrane, reflecting an *in vivo* PC distribution essential to the physiological action of PCs (**Fig. 2c-e**). More importantly, the CDDM differentiation protocol was highly reproducible when testing UB organoids derived from different genetic backgrounds (**Extended Data Fig. 3i**).

### Generating synthetic kidney from expandable NPCs and UBs

With the capacity to produce large quantities of NPCs and UPCs through *in vitro* expansion culture, we examined whether combining these cell types generated a model mimicking key features of *in vivo* kidney development, such as reiterative ureteric branching and nephron induction, and morphogenesis and patterning of differentiating derivatives (**Fig. 3a**).

**Figure 3.**
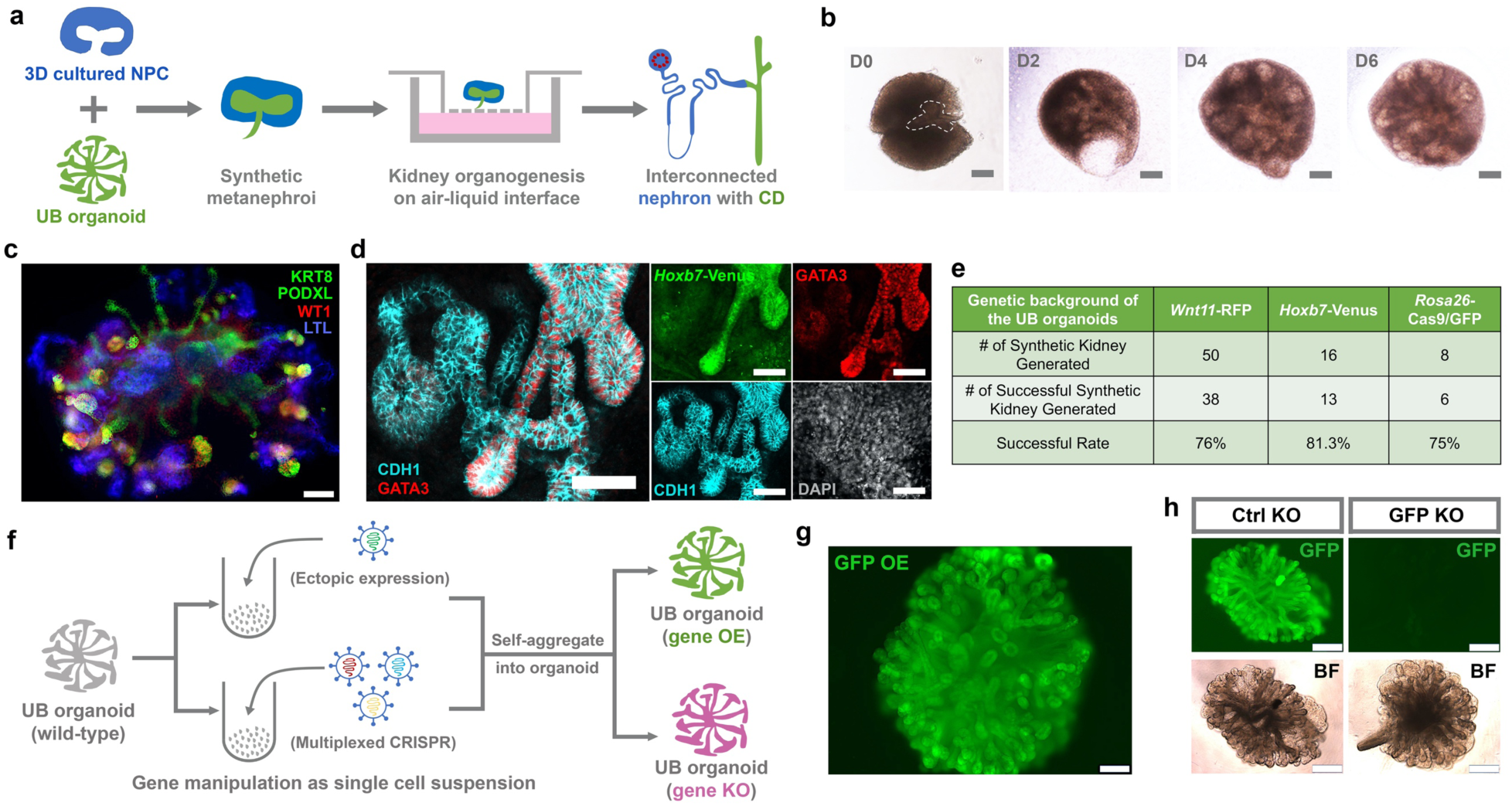
Synthetic kidney from expandable NPCs and UBs and gene editing of the UB organoid. **a**, Schematic of the synthetic kidney reconstruction and organotypic culture procedure. **b**, Time course images showing the branching morphogenesis of the synthetic kidney reconstruct on air-liquid interface. Dashed line indicates the UB organoid tip inserted into the excavated NPC aggregate. Scale bars, 100 µm. **c**, Immunostaining of the synthetic kidney (Day 7) constructed from *Wnt11*-RFP UB organoid and wild-type NPCs for UB/CD marker KRT8, nephron marker PODXL and WT1 (podocytes) and LTL (proximal tubule). Note that both KRT8 and PODXL were both stained green. The round structures that co-stain with WT1 are podocytes of the nephron. UB-derived structures do not co-stain with WT1. Scale bars, 100 µm. **d**, Immunostaining of the synthetic kidney constructed from *Hoxb7*-Venus UB organoid and wild-type NPC (Day 10) for GATA3, *Hoxb7*-Venus, and CDH1. Scale bars, 50 µm. **e**, Summary of synthetic kidney generation experiment. **f**, Schematic of gene overexpression and gene knockout procedures in UB organoid. OE, overexpression; KO, knockout. **g**, fluorescence image of GFP overexpression (GFP OE) in wild-type UB organoid. Scale bar, 200 µm. **h**, Knockout of GFP in *Rosa26*-Cas9/GFP UB organoid using multiplexed sgRNAs (“GFP KO”, right panels) targeting the coding sequence of GFP. Multiplexed non-targeting sgRNAs were introduced to the organoid as control (“Ctrl KO”, left panel). Note the gene-edited single cells self-organized into typical branching organoid morphology. Scale bars, 400 µm.

NPCs in our long-term culture model grow as 3D aggregates^14,25^. To mimic the natural organization of NPCs capping UB tips in the kidney anlagen, we manually excavated a hole in 3D cultured NPCs and inserted a cultured UB organoid tip into this cavity. The synthetic structures were then embedded into Matrigel and transferred onto an air-liquid interface (ALI) to facilitate further kidney organogenesis^45^. Over 7 days of culture, the UB underwent extensive branching (**Fig. 3b**) generating a KRT8^+^ tubular network extending from the center of the structure into the periphery. Further, NPCs generated nephron-like cell types and structures such as PODXL+/WT1+ podocytes and LTL+ proximal tubules (**Fig. 3c**). To determine whether the synthetic kidney also formed a connection between nephron and CD, we generated synthetic kidney from *Hoxb7*-Venus UB and wild-type NPCs. In this way, all progeny of the UB organoid could be tracked by Venus expression. Co-staining of the synthetic kidney structure with antibodies against CDH1 and GATA3 identified a clear fusion of CDH1^+^/Venus^-^ distal nephron with CDH1^+^/Venus^+^ CD. Importantly, GATA3 expression was strong in the entire Venus^+^ CD structure, but progressively dropped along the distal-to-proximal axis of the distal nephron, as observed *in vivo*^46,47^ (**Fig. 3d**). Thus, the synthetic kidney established an interconnection between the nephron and CD similar to that observed *in vivo*. Approximately 80% of synthetic kidneys underwent a similar program, indicating *in vitro* self-organizing development was robust (**Fig. 3e**). Most failures likely reflected technical differences in manual reconstruction. Taken together, synthetic kidney with interconnected nephron and CD can be efficiently generated from expandable NPCs and UBs.

### Performing efficient gene editing in the expandable UB organoid

The UB and CD models could provide an accessible *in vitro* complement to the mouse models for in-depth mechanistic studies and drug screening. Here, efficient gene overexpression (OE) or gene knockout (KO) would significantly extend the capability and utility of the *in vitro* model (**Fig. 3f**). As a proof of concept, GFP OE and GFP KO UB organoids were generated. For GFP OE, we used a standard lentiviral system to introduce GFP under the control of a CMV promoter^48^. However, even at a very high titer, the lentiviral infection efficiency of the intact T-shaped UB was very low (data not shown). But, dissociating T-shaped UB or UB organoids into a single cell suspension prior to infection, dramatically improved the infection efficiency, with widespread GFP activity in resulting UB organoids after re-aggregation of infected cells (**Fig. 3g**). To test gene-knockout, we targeted GFP in *Rosa26*-Cas9/GFP UB organoid, in which Cas9 and GFP are constitutively expressed from the *Rosa26* loci^34^ (**Extended Data Fig. 1e**). A mix of three different lentiviruses, expressing three different synthetic guide RNAs (sgRNAs) with Cas9 targeting sites 100-150bp apart^49^, gave a highly-efficient, GFP gRNA-specific, multiplexed CRISPR/Cas9 gene knockout, demonstrating effective gene knockout in UB organoid cultures (**Fig. 3h**).

### Generating human UB organoids from primary human UPCs

The successful generation of mouse UB and CD organoids prompted us to test whether the system can also derive human UB and CD organoids. To achieve this, we first developed a method to generate expandable human UB organoids from primary human UPCs (hUPCs) (**Extended Data Fig. 4a**). Similar to their murine counterparts, hUPCs within UB tips, express *RET*^*46,47,50*^ and *WNT11* (GUDMAP/RBK Resources, https://www.gudmap.org). Using an anti-RET antibody raised against the extracellular domain of RET that recognizes RET^+^ hUPCs (**Extended Data Fig. 4b**), we performed FACS enrichment of RET^+^ cells from human kidneys between 9-13 weeks of gestational age and examined their growth in modified UBCM conditions (**Extended Data Fig. 4c**). A robust human UBCM (hUBCM) culture condition was identified that sustained the long-term expansion (an estimated 10^8^-10^9^ fold expansion over 70 days) as branching UB organoids (**Fig. 4f; Extended Data Fig. 4d**). UPC marker gene expression was maintained in hUB cultures at level comparable to that of the human fetal kidney (**Extended Data Fig. 4e**).

**Figure 4.**
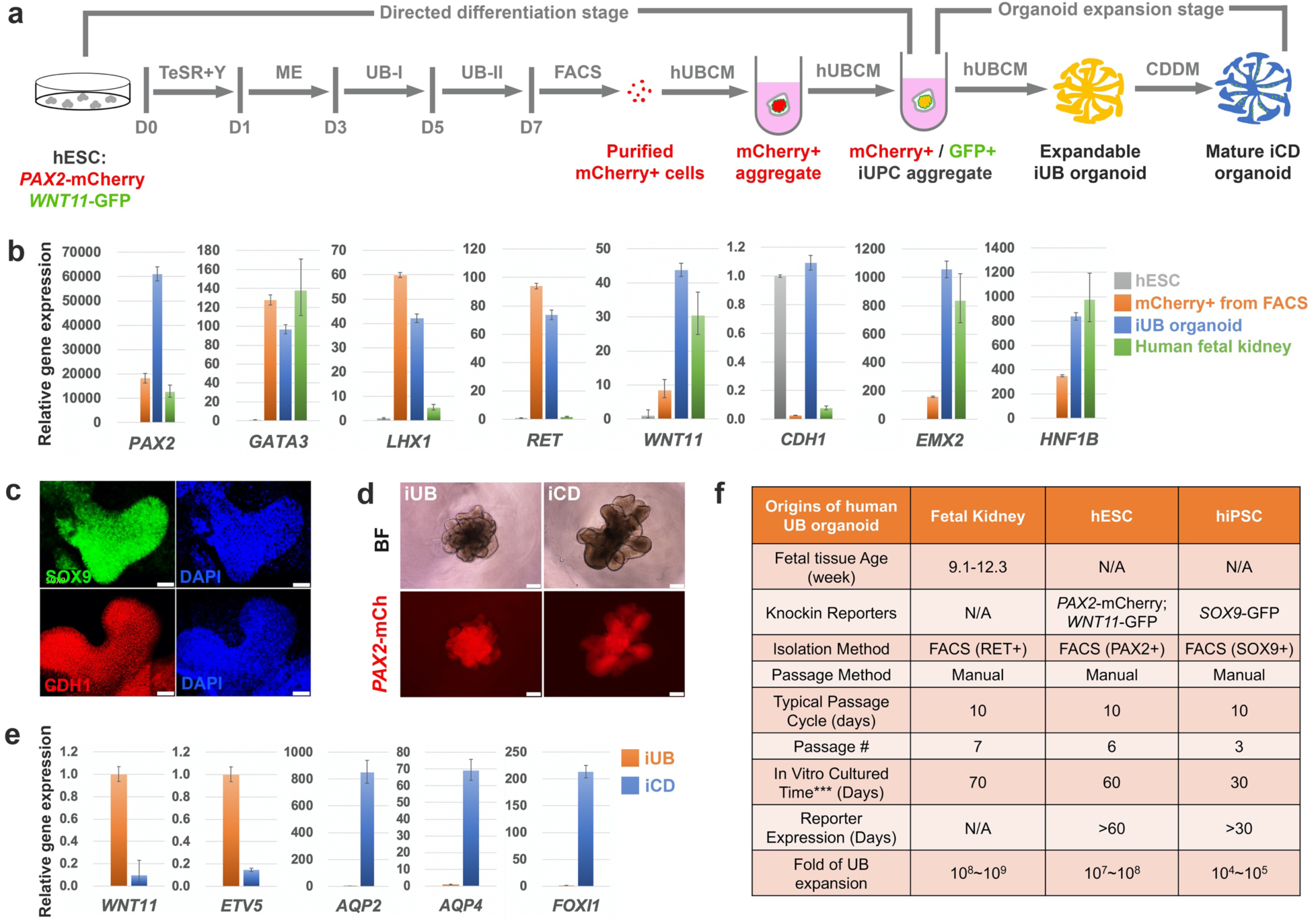
Generation of iUB and iCD organoids from human pluripotent stem cells. **a**, Schematic of the stepwise differentiation from *WNT11*-GFP/*PAX2*-mCherry dual reporter hESC line into iUB and iCD organoid. **b**, qRT-PCR analyses of the FACS purified mCherry+ cells (orange) and the iUB organoid (blue) for various UB lineage markers (*PAX2, GATA3, CDH1, HNF1B, LHX1, EMX2*), and UPC markers (*RET, WNT11*). Undifferentiated H1 hESCs (gray) and human fetal kidney (green, 11.2 week gestational age) were used as controls. Data are presented as mean ± s.d. Each column represents 3 technical replicates. **c**, Whole-mount immunostaining of the expandable iUB organoid for UB markers SOX9 and CDH1. Scale bars, 50 µm. **d**, Bright field (BF) and *PAX2*-mCherry (*PAX2*-mCh) images of expandable iUB organoid (left panels) and mature iCD organoid (right panels). Scale bars, 200 µm. **e**, qRT-PCR analyses of the iUB organoid (orange) and iUB-derived iCD organoid (blue), for UPC markers (*WNT11, ETV5*), PC markers (*AQP2, AQP4*), and IC marker (*FOXI1*). Data are presented as mean ± s.d. Each column represents 3 technical replicates. **f**, Summary of human UB organoid derivation from different sources and their expansion *in vitro*. ***, this is the culture time we achieved before our lab shutdown due to coronavirus outbreak. The maximum organoid culture time and expansion could be much greater.

### Generating iUB and iCD organoids from human pluripotent stem cells

To determine whether UB and CD organoids could be generated from human pluripotent stem cells (hPSC)-derived UPCs, we first genetically engineered H1 human embryonic stem cells (hESCs) with a knockin dual reporter system where mCherry was expressed from the *PAX2* locus (*PAX2*-mCherry) and GFP from the *WNT11* locus (*WNT11*-GFP) (**Extended Data Fig. 5**). This reporter cell line facilitated the establishment of a stepwise protocol that produced high-quality hUPCs, which generated branching induced UB (iUB) organoids, which matured into induced CD (iCD) organoids (**Fig. 4a**).

The UB is derived from the nephric duct (ND), which originates from primitive streak (mesendoderm)-derived anterior intermediate mesoderm^2,4^. Consistent with this developmental trajectory, following a 7-day directed differentiation protocol modified from previous protocols^22,23^, we were able to first observe the expression of mesendoderm (ME) marker T on day 3 of differentiation in most cells (**Extended Data Fig. 6a**), followed by the formation of large numbers of compact cell colonies that are GATA3^+^/SOX9^+^/PAX2^+^/PAX8^+^/KIT^+^/KRT8^+^ on day 7 of differentiation, suggesting the generation of potential precursor cells of the UB lineage (**Extended Data Fig. 6b**). Consistent with the immunostaining results, we were able to identify a *PAX2*-mCherry^+^ population (13.1%) by FACS on day 7. However, at this stage, the *PAX2*-mCherry^+^/*WNT11*-GFP^+^ population was very rare (0.4%), preventing further characterization or culture (**Extended Data Fig. 6c**). However, further culture of *PAX2*-mCherry+ cells in the 3D/hUBCM culture conditions activated *WNT11*-GFP reporter expression at around 3 weeks, and the structure started to show a branching morphology (**Extended Data Fig. 6d**). We refer to the *PAX2*-mCherry^+^/*WNT11*-GFP^+^ branching structure an “iUB” organoid hereafter. Importantly, these iUB organoids could be expanded stably in 3D/hUBCM for at least 2 months without losing reporter gene expression (**Fig. 4f**). Consistently, qRT-PCR analysis confirmed that *WNT11* expression was low in the mCherry^+^ cells purified from FACS, but was dramatically elevated in the iUB organoid. Furthermore, even though UB marker genes *PAX2, GATA3, LHX1* and *RET* were greatly elevated on day 7 of differentiation, while *WNT11, CDH1, EMX2*, and *HNF1B*, showed comparable levels of expression to the human fetal kidney only after extended hUBCM culture, suggesting hUBCM promoted transition from a common nephric duct to a specific ureteric epithelial precursor (**Fig. 4b**). In addition to qRT-PCR, expression of marker genes *SOX9* and *CDH1* in the iUB organoid was detected at the protein level by immunostaining (**Fig. 4c**), further confirming the identity of the iUB organoid.

We next tested whether the expandable iUB organoid retained the potential to generate an iCD organoid after long-term expansion. iUB organoids were subjected to differentiation with the CDDM medium identified for mouse UB-to-CD transition. After 14 days of differentiation in CDDM, the human iUB organoid grew and elongated, maintaining *PAX2*-mCherry expression, but losing *WNT11*-GFP expression (**Fig. 4d**, and data not shown). More importantly, the expression of UPC markers *WNT11* and *RET* was greatly diminished, while CD marker genes *AQP2, AQP4* and *FOXI1* were dramatically elevated, suggesting the successful transition from iUB to iCD (**Fig. 4e**).

Lastly, we tested whether expandable iUB organoids could be generated from human induced pluripotent stem cells (hiPSCs). For this, we employed *SOX9*-GFP hiPSC^51^ for differentiation and purified the *SOX9*-GFP+ UB precursor cells on day 7 of differentiation (**Extended Data Fig. 6e**). Similar to hESC-derived iUBs, following an extended culture in hUBCM, we were able to derive *SOX9*-GFP iUB organoids that expanded stably with retained *SOX9*-GFP expression throughout (**Fig. 4f; Extended Data Fig. 6f-h**). Taken together, these results support the conclusion that expandable iUBs and mature iCDs organoid can be derived from hESC and hiPSC lines.

## DISCUSSION

Here, we show that with appropriate 3D culture conditions, it is possible to expand UB progenitor cells *in vitro* as branching UB organoids. The organoid culture medium thus serves as a synthetic niche for maintaining the UB progenitor cell identity. Consistent with mouse genetics studies, signaling pathways that play key roles in kidney branching morphogenesis, such as GDNF^27,52-55^, FGF^56^, RA^57-59^ and Wnt^60,61^ signaling, are also essential in maintaining UB progenitor cell identity in UB organoids. Combined with efficient genome editing, this accessible *in vitro* model for UB progenitor cell self-renewal is a powerful tool for comparative studies of how UB progenitor cell fate is determined in mouse and human.

Leveraging our ability to produce large quantities of high-quality UB progenitor cells in the format of expandable branching UB organoids, we performed a screening that identified CDDM— a cocktail of growth factors, small molecules and hormones that together can differentiate UB organoids into CD organoids with spatially patterned mature PCs and ICs. The molecular mechanisms underlying the UB-to-CD transition are still largely unknown. The *in vitro* organoid system provides a novel tool to study this process, and the chemically-defined components in CDDM shed new light on potential signals that trigger CD maturation *in vivo*. Despite the general difficulty of maturing stem cell-derived tissues, our study shows that it is possible to achieve proper patterning and maturation *in vitro*, similar to what we observe *in vivo*, when starting from high-quality progenitor cells under appropriate culture conditions.

The generation of a synthetic kidney from expandable NPCs and UPCs provides a proof-of-concept for rebuilding a kidney *in vitro* from kidney-specific progenitor cells. The availability of expandable NPCs and UPCs provides the building blocks required for making a kidney. The interaction between NPCs and UPCs is faithfully recapitulated, leading to the autonomous differentiation into interconnected nephron and collecting duct structures. However, there are still many obstacles to generating a functional synthetic kidney for patients with kidney failure. For example, to generate a better organized kidney structure, the IPC population might be needed^22^, as well as the vascular progenitor population. To generate these populations, we might develop strategies similar to those used to produce NPCs and UPCs via primary progenitor cell expansion or hPSC differentiation. In addition, transplantation of the immature synthetic metanephroi *in vivo* might facilitate the generation of vasculature^14,62,63^. Furthermore, we need engineering methods for scaling up the production of nephrons and CDs in order to provide meaningful filtration capacity in a synthetic kidney transplant. Nevertheless, the availability of expandable NPCs and UPCs lays the foundation for achieving this ambitious goal in the future.

Efficient genome editing in UB organoids opens up many new applications using the UB and CD organoid platform. UB and CD organoids can be generated from available transgenic mouse strains that bear genetic mutations related to kidney development and disease. In addition, disease-relevant mutations can be introduced into the UB organoid directly, enabling the investigation of pathophysiology throughout the entire course of kidney branching morphogenesis, from the UB branching period to the mature CD stage. The ability to produce large quantities of UB and CD organoids also provides a platform for drug screening. The mature PCs and ICs present in the CD organoids are potential sources for cell replacement therapies for patients with CD damage. In conclusion, the novel UB and CD organoid system provides a powerful tool for studying kidney development, modeling kidney disease, discovering new drugs and, ultimately, regenerating the kidney.

## Acknowledgments

We would like to thank Jeffrey Boyd and Bernadette Masinsin of the USC Flow Cytometry Facility for FACS, Seth Ruffins of the USC Optical Imaging Facility for help with microscopy, Dejerianne Ostrow and David Ruble of the Children’s Hospital Los Angeles Sequencing Core for RNA-seq, Meng Li and Yibu Chen of the USC Norris Medical Library Bioinformatics Service for help with the RNA-seq computational analysis, Haruhiko Akiyama and Juan Carlos Izpisua Belmonte for sharing the *Sox9*-GFP mice, Naoki Nakayama for sharing the *SOX9*-GFP hiPSC line, Dr. Melissa L. Wilson (Department of Preventive Medicine, University of Southern California) and Family Planning Associates for coordinating fetal tissue collection, and Cristy Lytal for help with editing the manuscript.

## Author contributions

Z.Z., A.P.M., and Z.L. designed the study. Z.Z., B.H., R.K.P., Y.L., J.C., T.P., E.A.R., A.D.K. performed experiments. A.V. and M.E.T. performed RNA-seq computational analysis. M.E.T and B.H.G. provided human fetal kidney samples. N.P.S., K.R.H., and A.P.M. provided reagents and helpful discussions. Z.Z., B.H., J.Y., A.P.M., and Z.L. wrote the manuscript.

## Data and Materials availability

All data is available in the main text or the supplementary materials. RNA-seq data have been submitted to Gene Expression Omnibus (GEO) with accession number GSE149109.

## Supplementary Information

**Inventory of Supplementary Information**

I Methods

II Extended Data Figures 1-6

III Extended Data Tables 1-4

IV Culture Medium Recipes 1-6

V Supplemental References

## Methods

### Animal work

All animal work was performed under Institutional Animal Care and Use Committee approval (USC IACUC Protocol # 20829).

Swiss Webster mice were purchased from Taconic Biosciences (Model # SW-F, MPF 4 weeks).

*Wnt11-*RFP mice were kindly shared from Dr. Andrew McMahon (JAX # 018683).

*Sox9*-GFP mice were kindly shared from Dr. Haruhiko Akiyama^1^.

*Hoxb7*-Venus mice were kindly shared from Dr. Andrew McMahon (JAX # 016252).

*Rosa26-*Cas9/GFP mice were kindly shared from Dr. Andrew McMahon (JAX #026179).

### Derivation of mouse UB Organoid

#### From intact T-shaped UB

Male mice with desired genotype (*Wnt11*-RFP, *Hoxb7*-Venus, *Sox9-*GFP, or *Rosa26*-Cas9/GFP) were mated with female Swiss Webster mice. Plugs were checked the next morning, midday of plug positive was designated as embryonic day 0.5 (E0.5). Timed pregnant mice were euthanized at E11.5. Kidneys were dissected out from embryos with standard dissection technique and transferred into kidney dissection medium in a 1.5mL Eppendorf tube on ice. Next, the medium in the tube was removed, at least 500µL fresh pre-warmed collagenase IV was added into the tube and incubated at 37°C for 20-22 minutes. After incubation, collagenase was removed and at least 500µl mouse UB dissection medium was added to resuspend the kidneys. 1-3 kidneys were transferred each time with 40-80µL medium onto a 100mm petri dish lid as a working droplet. UBs were isolated from the surrounding MM and other tissues using sterile needles (BD, Cat. No. BD305106) without damaging UBs. The isolated UBs can be temporarily left in the medium at room temperature for less than 30 minutes while dissecting other UBs. After all UBs were isolated, each UB was transferred into an 8µl cold Matrigel droplet in one well of a U-bottom 96-well low-attachment plate, by using a P10 micropipette. The UB and Matrigel drop were mixed by pipetting up and down gently 2-3 times. After all UBs were embedded in Matrigel, the plate was incubated at 37°C for 20 minutes for the Matrigel to solidify. Then, 100µl mouse UBCM (mUBCM) was slowly added into each well and the plate was then transferred into an incubator set at 37°C with 5% CO_2_.

#### From dissociated UB single cells

For deriving UB organoid from dissociated UB single cells (e.g. for gene editing purpose), after the isolation of E11.5 T-shaped UBs from kidneys following the procedures described above, all UBs were collected into a 1.5mL Eppendorf tube with the medium removed as much as possible. Appropriate amount (e.g. for 20 UBs, we use 200µl, adjust accordingly) of pre-warmed Accumax cell dissociation solution (Innovative Cell Technologies, # AM105) was added into the tube and the tube was then incubated at 37°C for 25 minutes. Then, an equal amount of mouse UB dissection medium was added into the tube to neutralize the Accumax and the mixture was pipetted up and down gently for 25 times to dissociate the UB into single cells. The tube was then centrifuged at 300g for 5 minutes. After centrifugation, supernatant was carefully removed and UB cells were resuspended in appropriate amount of mUBCM (Y27632 was supplemented at 10µM final concentration for the first 24 hours) by pipetting up and down gently for less than 10 times. Cell density was measured using automatic cell counter (Bio-Rad, TC20). 2000 cells (∼1x E11.5 UB) were transferred into each well of a U-bottom 96-well low-attachment plate and extra amount of mUBCM (with 10µM Y27632) was added to the well to make the final volume 100µl per well. The plate was then centrifuged at 300g for 5 minutes and transferred and cultured in 37°C incubator. After 24 hours, the 2000 single cells formed an aggregate autonomously and the aggregate was then transferred into an 8µl cold Matrigel droplet in another well of the U-bottom 96-well low-attachment plate using a P10 micropipette. The aggregate was pipetted up and down gently for 2-3 times to mix with Matrigel. After all aggregates were embedded in Matrigel, the plate was incubated at 37°C for 20 minutes for the Matrigel to solidify. Lastly, 100µl UBCM was added slowly into each well and the plate was then transferred into an incubator set at 37°C with 5% CO_2_.

### Mouse UB organoid expansion and passaging

Mouse UBCM was renewed with fresh medium every two days, and UB organoid was passaged every five days.

#### For manual passaging as small tips

UB organoid (with Matrigel) was first transferred from U-bottom 96-well plate onto a 100mm petri dish lid with 80-100µl medium using a P1000 pipette with the tip cut 0.5-1 cm to widen the diameter. Most of the Matrigel surrounding the organoid was removed using sterile needles under dissecting microscope. A small piece of the organoid with 3-5 branching tips was cut using the needles and then transferred into an 8µl cold Matrigel droplet in a U-bottom 96-well low-attachment plate well by using a P10 micropipette. The small piece of tip structure was pipetted up and down 2-3 times to mix with the Matrigel. The plate was then incubated at 37°C for 20 minutes for the Matrigel to solidify. Then, 100µl mUBCM was slowly added into each well and the UB was cultured in 37°C incubator.

#### For passaging as single cells

UB organoid was transferred into a 1.5ml Eppendorf tube with as little medium as possible using a P1000 pipette with the tip cut 0.5-1 cm to widen the diameter. 200µl pre-warmed Accumax cell dissociation solution was added into the tube and gently pipetted up and down using a P200 pipette to break down the organoid (around 20 times) into small pieces. The tube was then incubated in 37°C for 25 minutes. After the incubation, 200µl UB dissection medium was added to neutralize the Accumax and the mixture was pipetted up and down gently for 25 times to further break down the organoid into single cells. The tube was then centrifuged at 300g for 5 minutes. After centrifugation, supernatant was carefully removed and the UB cells were resuspended in appropriate amount of mUBCM (with the addition of Y27632 at 10µM final concentration for the first 24 hours) by pipetting up and down gently for less than 10 times. Cell density was measured by automatic cell counter. 2000 cells were transferred into each well of a U-bottom 96-well low-attachment plate and extra amount of mUBCM (with 10µM Y27632) was added to the well to make the final volume 100µl per well. The plate was then centrifuged at 300g for 5 minutes and transferred and cultured in 37°C incubator. After 24 hours, the 2000 single cells formed an aggregate autonomously and the aggregate was then transferred into an 8µl cold Matrigel droplet in another well of the U-bottom 96-well low-attachment plate using a P10 micropipette. The aggregate was pipetted up and down gently for 2-3 times to mix with Matrigel. After all aggregates were embedded in Matrigel, the plate was incubated at 37°C for 20 minutes for the Matrigel to solidify. Lastly, 100µl UBCM was added slowly into each well and the plate was then transferred into an incubator set at 37°C with 5% CO_2_.

### Derivation of human UB organoid

#### Organoid derivation from RET+ primary UPCs purified from human fetal kidney

All human fetal kidney samples were collected under Institutional Review Board approval (USC-HS-13-0399 and CHLA-14-2211). Following the patient decision for pregnancy termination, the patient was offered the option of donation of the products of conception for research purposes, and those that agreed signed an informed consent. This did not alter the choice of termination procedure, and the products of conception from those that declined participation were disposed of in a standard fashion. The only information collected was gestational age and whether there were any known genetic or structural abnormalities. The kidney nephrogenic zone was dissected manually from fresh 9-13-week human fetal kidney, chopped into small pieces with surgical blade, and transferred into 4-6 1.5ml Eppendorf tubes. Tissues were washed with PBS and resuspended with 500µl of pre-warmed Accumax per tube and the tubes were incubated at 37°C with shaking for 25 minutes. 500µl dissection medium was then added to neutralize the Accumax, and the mixture was pipetted up and down for 25 times to dissociate the tissues into single cells. The mixture medium with kidney cells were then combined together and went through a 40µm cell strainer, then transferred into 1.5ml Eppendorf tubes. These tubes were then centrifuged at 300g for 5 minutes and the supernatant was carefully removed. The kidneys cells in each tube were then resuspended and combined in total 300-400µl cold FACS medium (1x PBS, 1X Pen Strep, 2% FBS) with human anti-RET antibody at 1:200 dilution into one tube and incubated for 30 minutes on ice. The tube was gently tapped every 10 minutes. After 30 minutes, 1ml cold FACS medium was added into the tube. The tube was then centrifuged at 300g for 5 minutes and the supernatant was carefully removed. Cells in the tube were resuspended again with 500µl cold FACS medium plus secondary antibody (Donkey anti-Goat, Alexa Fluor 568, Invitrogen, Cat. # A-11057) at 1:1000 and incubated for 30 minutes on ice. The tube was gently tapped every 10 minutes. After the incubation, 1ml cold FACS medium was added into the tube. The tube was then centrifuged at 300g for 5 minutes and the supernatant was carefully removed. Cells in the tube were resuspended with 300-500µl cold FACS medium plus DAPI at 1:2000 ratio, went through 40µm cell strainer and transferred into a FACS tube, and placed on ice before FACS. RET^+^ (Alexa Fluro 568) UPCs were then sorted out by FACS. The RET+ cells were collected in a 1.5ml Eppendorf tube with 500µl dissection medium. The tube was centrifuged at 300g for 5 minutes and the supernatant was carefully removed. These cells were then resuspended in appropriate amount of hUBCM. Cell density was measured by automatic cell counter. 2000-20000 cells were transferred into each well of a U-bottom 96-well low-attachment plate and appropriate amount of hUBCM was added to the well to make the final volume to be 100µl per well. After 24 hours, UB cell aggregate was formed and transferred into an 8µl cold Matrigel droplet in another well of the U-bottom 96-well low-attachment plate by using a P10 micropipette. The aggregate was pipetted up and down 2-3 times gently to mix with the Matrigel. After all aggregates were embedded in Matrigel, the plate was incubated at 37°C for 20 minutes for the Matrigel to solidify. Lastly, 100µl hUBCM was added slowly into each well and cultured in 37°C incubator. After around 10-15 days of culturing, epithelial tip structures can be seen budding out from the aggregate. These tip structures were dissected out and re-embedded into Matrigel and expanded as human UB organoid.

#### Organoid derivation from human ESCs and iPSCs

Human pluripotent stem cells are routinely cultured in mTeSR1 (TeSR) medium in monolayer culture format coated with Matrigel and passaged using dispase as previously described^2^. The hPSCs were pre-treated with 10µM Y27632 in TeSR medium for one hour before dissociated into single cells using Accumax cell dissociation solution. Following dissociation, 60,000 cells were seeded into Matrigel coated 12-well plate with 1ml TeSR medium plus 10µM Y27632 (day 0). 24 hours later (day 1), the medium was removed and 1ml ME stage medium was slowly added to the well. 48 hours later (day 3), ME stage medium was removed and 1ml UB-I stage medium was slowly added to the well. 24 hours later (day 4), medium was changed to 1ml fresh UB-I medium again. After another 24 hours (day 5), UB-I medium was removed, and 1.5-2ml UB-II stage medium was slowly added to the well. 24 hours later (day 6), medium was changed to 1.5-2ml fresh UB-II medium. At day 7, differentiated cells were dissociated into single cells following standard Accumax dissociation method. Dissociated cells were resuspended in 300-500µl cold FACS medium plus DAPI at 1:2000 ratio, went through 40µm cell strainer (Greiner bio-one, Cat. No. 542040), and placed on ice before FACS. mCherry^+^ cells (from *PAX2*-mCherry/*WNT11*-GFP hESC line) or GFP^+^ cells (from *SOX9*-GFP hiPSC line) were then sorted out by FACS. Upon FACS sorting of the mCherry^+^ cells or GFP^+^ cells, these cells were collected in a 1.5ml Eppendorf tube with 500µl dissection medium. The tube was centrifuged at 300g for 5 minutes and the supernatant was carefully removed. These cells were then resuspended with appropriate amount of hUBCM. 2000-20000 cells were transferred into each well of a U-bottom 96-well low-attachment plate and appropriate amount of hUBCM was added to the well to make the final volume to be 100µl per well. After 24 hours, UB cell aggregate was formed and transferred into an 8µl cold Matrigel droplet in another well of the U-bottom 96-well low-attachment plate by using a P10 micropipette. The aggregate was pipetted up and down 2-3 times gently to mix with the Matrigel. After all aggregates were embedded in Matrigel, the plate was incubated at 37°C for 20 minutes for the Matrigel to solidify. Lastly, 100µl hUBCM was added slowly into each well and cultured in 37°C incubator. After around 10-15 days of culturing, epithelial tip structures can be seen budding out from the aggregate. These tip structures were dissected out and re-embedded into Matrigel to continue culture. *WNT11*-GFP expression was induced within another week (totaling 25 days post FACS on average). From then, the iUB organoid was established and can be passaged stably following the procedures below.

### Human UB organoid expansion and passaging

Both human UB organoid from primary RET^+^ UPC and iUB organoid from hPSCs were passaged every 9-11 days. The passaging methods (manual or as single cells) are the same as passaging mouse UB organoid, with the exception of using hUBCM instead of mUBCM.

### CD Differentiation

#### Mouse CD differentiation

Mouse UB organoid was passaged at day 5 of expansion as single cells and 1700 cells were seeded for continuing expansion. At day 10 of mUB expansion, mUBCM was removed and 150µl 1xPBS was added and removed to wash the organoid. 100µl CD differentiation medium (CDDM) was then added to start mouse CD (mCD) differentiation (mCD differentiation Day 0). Organoid was cultured in 37°C incubator and medium was changed every two days or everyday if needed (when organoid is big, and the medium turns yellow within 24 hours) for a total of seven days. No passage is needed. At mCD differentiation Day 7, mCD organoid was harvest for analyses.

#### Human CD differentiation

After human UB organoid expansion was stabilized (at least 25 days post FACS), hUBCM was removed and 100µl CD differentiation medium was added to start hCD differentiation (hCD differentiation Day 0). Organoid was cultured in 37°C incubator and medium was changed every two days or everyday if needed (when organoid is big, and the medium turns yellow within 24 hours) for a total of 14 days. Organoid can be passaged if it is too big, but it is better to start with smaller organoid. At hCD differentiation Day 14, hCD organoid was harvest for analyses.

### Mouse Synthetic Kidney Generation

A small piece (with 2-4 branching tips) of Day 7-9 cultured mUB organoid was inserted into a microdissected hole on a 3D cultured mNPC aggregate in kidney reconstruction medium (APEL2 + 0.1µM TTNPB) with 10µM Y27632. The reconstruct was then transferred into a well of a U-bottom 96-well low-attachment plate with 100µl kidney reconstruction medium plus 10µM Y27632, using a P1000 pipette with the top 0.5-1 cm of the tip cut, and cultured in 37°C incubator (day 0). After 24 hours (day 1), dead cells surrounding the reconstruct was removed by pipetted up and down several times in the well using a P1000 pipette with the top 0.5-1 cm of the tip cut. The reconstruct was then transferred and embedded into 10µl Matrigel droplet in another well using the same P1000 pipette and tip or a smaller pipette and tip if possible. Sterile needle can be used to position the reconstruct in Matrigel droplet. The plate was then incubated at 37°C for 20 minutes for the Matrigel to solidify. Then, 100µl kidney reconstruction medium was slowly added into the well and the reconstruct was cultured in 37°C incubator. After another 24 hours (day 2), the reconstruct was transferred together with its surrounding Matrigel onto a 12-well transwell insert membrane using a P1000 pipette with the top 0.5-1 cm of the tip cut. 300µl kidney reconstruction medium was added in the lower chamber of the transwell. The medium was changed every two days for a total of 10-12 days for the kidney reconstruction, then the reconstruct was harvested for analyses.

### FACS

Cells were dissociated/prepared as described above. FACS sorting was performed on a BD FACS ARIA lllu cell sorter. Sorted cells were collected in a 1.5ml Eppendorf tube with 500µl dissection medium on ice.

### RNA isolation, reverse transcription and quantitative PCR

Samples were dissolved in 100µl TRIzol (Invitrogen, Cat. No. 15596018) and kept in −80°C freezer. RNA isolation was performed using the Direct-zol RNA MicroPrep Kit (Zymo Research, Cat. No. R2062) according to the manufacturer’s instructions. Reverse transcription was performed using the iScript Reverse Transcription Supermix (Bio-Rad, Cat. No. 1708841) following the manufacturer’s instructions. qRT-PCR was performed using the Applied Biosystems PowerUp SYBR Green Master Mix (Thermo Fisher, Cat. No. A25777) and carried out on an Applied Biosystems Vii 7 RT-PCR system (*Life Technologies*). Validated gene-specific primers can be found in Extended Data Table 2. Fold change was calculated from ΔCt using *Gapdh* as housekeeping gene as previously described^3^.

### Immunofluorescence

#### Whole-mount staining

Samples were fixed in 4% PFA for 5 minutes (UB/CD organoid) or 20 minutes (kidney reconstruct) in Eppendorf tubes or tissue culture plates at room temperature. They were then washed four times in 1X PBS (Corning, Cat. No. 21-040-CV) for total 30 minutes. After wash, samples were blocked in blocking solution (0.1% PBST containing 3% BSA) for 30 minutes at room temperature or 4°C overnight followed by primary antibody staining (primary antibodies were diluted in blocking solution) at 4°C overnight. On the second day, samples were washed four times with 0.1% PBST for total 30 minutes at room temperature. Secondary antibodies diluted in blocking solution were added and samples were incubated at 4°C overnight. On the third day, samples were washed four times with PBST for total 30 minutes at room temperature.

#### Cryo-section staining

Samples were fixed in 4% PFA for 5 minutes (UB/CD organoid) in Eppendorf tubes or tissue culture plates at room temperature and then washed four times in 1X PBS for total 30 minutes. They were then transferred into a plastic mold and embedded in OCT Compound (Scigen, Cat. No. 4586K1) and froze in −80°C for 24 hours to make a cryo-block. The cryo-blocks were sectioned using Leica CM1800 Cryostat. The sectioned slides were then blocked for 30 minutes at room temperature followed by one hour of primary antibodies staining at room temperature. The slides were then washed four times with PBST for five minutes, and then secondary staining for 30 minutes. After the secondary staining, the slides were washed four times with PBST for five minutes and mounted with mounting medium.

### *WNT11-*GFP/*PAX2-*mCherry dual reporter hESC line generation

CRISPR-Cas9 based genome editing was used to insert 2A-EGFP-FRT-PGK-Neo-FRT or 2A-mCherry-loxP-PGK-Neo-loxP cassette downstream of the stop codon (removed) of endogenous *WNT11* or *PAX2* gene, respectively. DNA sequences ∼1Kb upstream and ∼1K downstream of endogenous *WNT11* (upstream F: CCGGAATTCGACGTAATCATTCCACTGACC; upstream R: TACGAGCTCCTTGCAGACATAGCGCTCCAC; downstream F: CGCGTCGACGGCCCTGCCCTACGCCCCA; downstream R: CCCAAGCTTTGCCTGGAAACTGGAGAGCTCCCTC) and *PAX2* (upstream F: GAAGTCGACTTTCCACCCATTAGGGGCCA; upstream R: TATGCTAGCGTGGCGGTCATAGGCAGCGG; downstream F: TATACGCGTTTACCGCGGGGACCACATCA; downstream R: GACGGTACCAGTAACTGCTGGAGGAAGAC) stop codon were cloned upstream and downstream of 2A-EGFP-FRT-PGK-Neo-FRT or 2A-mCherry-loxP-PGK-Neo-loxP cassette respectively to facilitate homologous recombination. 2A-EGFP fragment was cloned from pCAS9_GFP (Addgene #44719) and the FRT-PGK-Neo-FRT cassette was cloned from pZero-FRT-Neo3R (kindly provided by Dr. Keiichiro Suzuki). 2A-mCherry-loxP-Neo-loxP fragment was cloned from Nanog-2A-mCherry plasmid (Addgene #59995). The different fragments were then cloned to pUC19 plasmid to make the complete donor plasmids for both knockin experiments. gRNA oligos for WNT11 (F: CACCGGTCCTCGCTCCTGCGTGGGG; R: AAACCCCCACGCAGGAGCGAGGACC) and PAX2 (F: CACCGATGACCGCCACTAGTTACCG; R: AAACCGGTAACTAGTGGCGGTCATC) were synthesized and cloned into the lentiCRISPR v2 plasmid (Addgene # 52961). First, both donor and gRNA plasmids for *PAX2* reporter KI were transfected into the H1 hESCs using the Lipofectamine 3000 Transfection Reagent (Invitrogen, Cat. No. L3000015). Neomycin-resistant single cell colonies were picked up manually and genotyping was performed based on PCR. PCR primers see Extended Data Table 1, results see Extended Data Fig. 5. Clones with biallelic knockin of *PAX2*-mCherry were chosen for second round screen where plasmid encoding Cre was delivered to allow the transient expression Cre, whose activities excise the loxP-flanked PGK-Neo cassette from the knockin alleles. PCR was performed to identify the single cell clones in which PGK-Neo cassettes were excised from both alleles. Then the same strategy was used to knock in *WNT11* reporter based on the successful biallelic *PAX2*-mCherry knockin clones.

### RNA-seq and data analysis

Day 5 and day 10 cultured mUB organoids were collected and prepared for RNA-Seq. RNA sequencing was analyzed using Partek Flow, including published dataset of interstitial progenitor cells and nephron progenitor cells (Lindstrom et al., 2018), ureteric tip and trunk cells (Rutledge, et al., 2017). FASTQ files were trimmed from both ends based on a minimum read length of 25bps and a shred quality score of 20 or higher. Reads were aligned to GENCODE mm10 (release M23) using STAR 2.6.1d. Aligned reads were quantified to the Partek E/M annotation model. Gene counts were normalized by adding 1 then by TMM values. Samples were filtered to include differentially expressed genes of UB Tip compared to UB Trunk with false discovery rate <= 0.01 and fold change <-5 or >5, resulting in 1841 UB Tip signature genes. Then, hierarchical clustering was produced on by clustering samples and features with average linkage cluster distance and Euclidean point distance. Principle component analysis (PCA) was performed using the EDASeq R/Bioconductor packages and the plot was rendered with the ggplot2 R package.

### Gene editing in mUB organoid

#### Gene over-expression

Lentiviral infection was used to overexpress GFP in E11.5 mUB cells. Lentivirus was first concentrated 100x using Lenti-X Concentrator kit from Takara (Cat # 631231). Concentrated lentivirus was aliquoted and stored in −80 before use. The lentivirus was used at 1x final concentration together with 10μM Polybrene (Sigma-Aldrich, Cat. No. TR-1003-G) diluted in mUBCM. 100µl virus-UBCM mixture was added to the U-bottom 96-well low-attachment plate well with single cells suspension prepared from 8-10 E11.5 mUBs. The UBs and virus were centrifuged together at 800g for 30 minutes for spinfection^4^ at room temperature. After the spinfection, virus-UBCM mixture was removed and the infected UB cells were washed three times with PBS, then aggregated and embedded in Matrigel and cultured in mUBCM in 37°C incubator following standard UB organoid culture procedures described above. 200μg/ml G-418 (Invitrogen, Cat. # 10131027) was applied to select for UB cells that have been successfully infected. The UB aggregate self-organize into typical branching organoid 4-5 days after infection.

**Extended Data Fig. 1.**
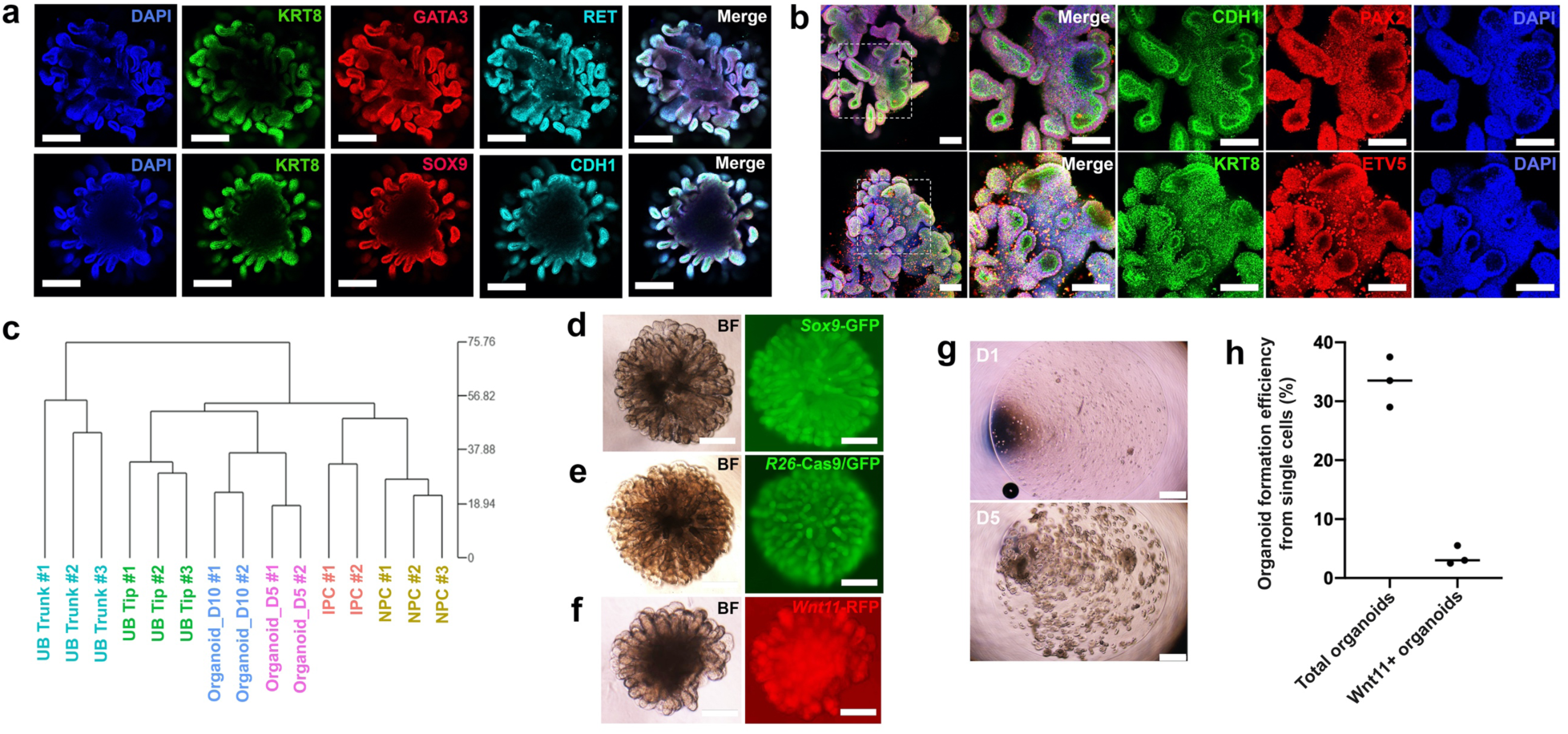
Derivation and characterization of mouse UB organoid. **a, b**, Immunostaining of the branching UB organoid after 10 days of culture for various UB markers. Note the red signals that do not overlay with DAPI from PAX2 and ETV5 staining panels are non-specific signals. Scale bars, **a**, 200 µm; **b**, 100 µm. **c**, Unsupervised clustering analysis of RNA-seq data. **d**, Bright field (BF) and fluorescence images of UB organoid derived from *Sox9*-GFP genetic background. Scale bars, 200 µm. **e**, Bright field (BF) and fluorescence images of UB organoid derived from *Rosa26*-Cas9/GFP (*R26*-Cas9/GFP) genetic background. Scale bars, 200 µm. **f**, Bright field (BF) and fluorescence images of *Wnt11*-RFP UB organoid revived from freezing. Scale bar, 200 µm. **g**, Bright field images showing 200 single cells embedded into one drop of Matrigel and cultured in UBCM for 1 day (D1) and 5 days (D5). Scale bars, 500 µm. **h**, Efficiency of UB organoid formation from 200 single UB cells. Data are presented as mean ± s.d. Each group represents 3 biological replicates.

**Extended Data Fig. 2.**
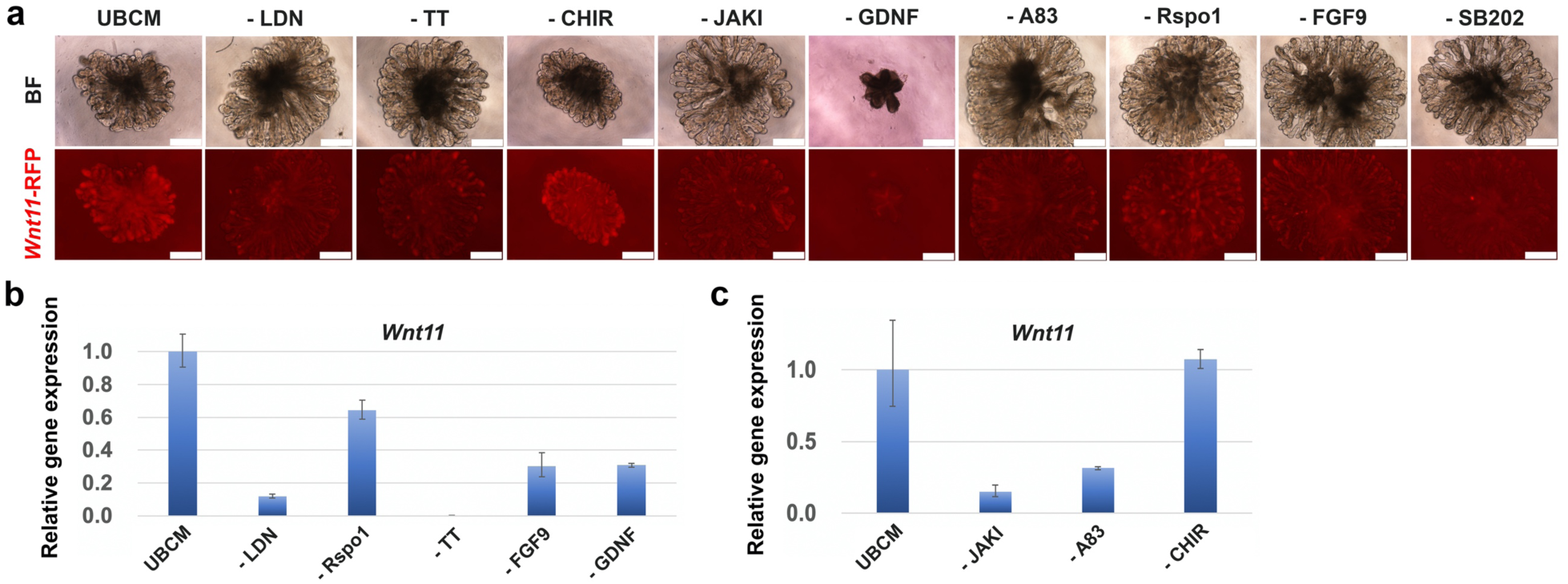
UBCM represents a synthetic niche for UPC self-renewal. **a**, Bright field (BF) and fluorescence images showing the morphology and *Wnt11*-RFP reporter expression in the UB organoids cultured in complete UBCM or upon withdrawal of indicated components from the UBCM. Scale bars, 200 µm. **b, c**, qRT-PCR analyses of the UB organoids from (**a**) for the UB progenitor marker gene *Wnt11*. Data are presented as mean ± s.d. Each column represents 3 technical replicates.

**Extended Data Fig. 3.**
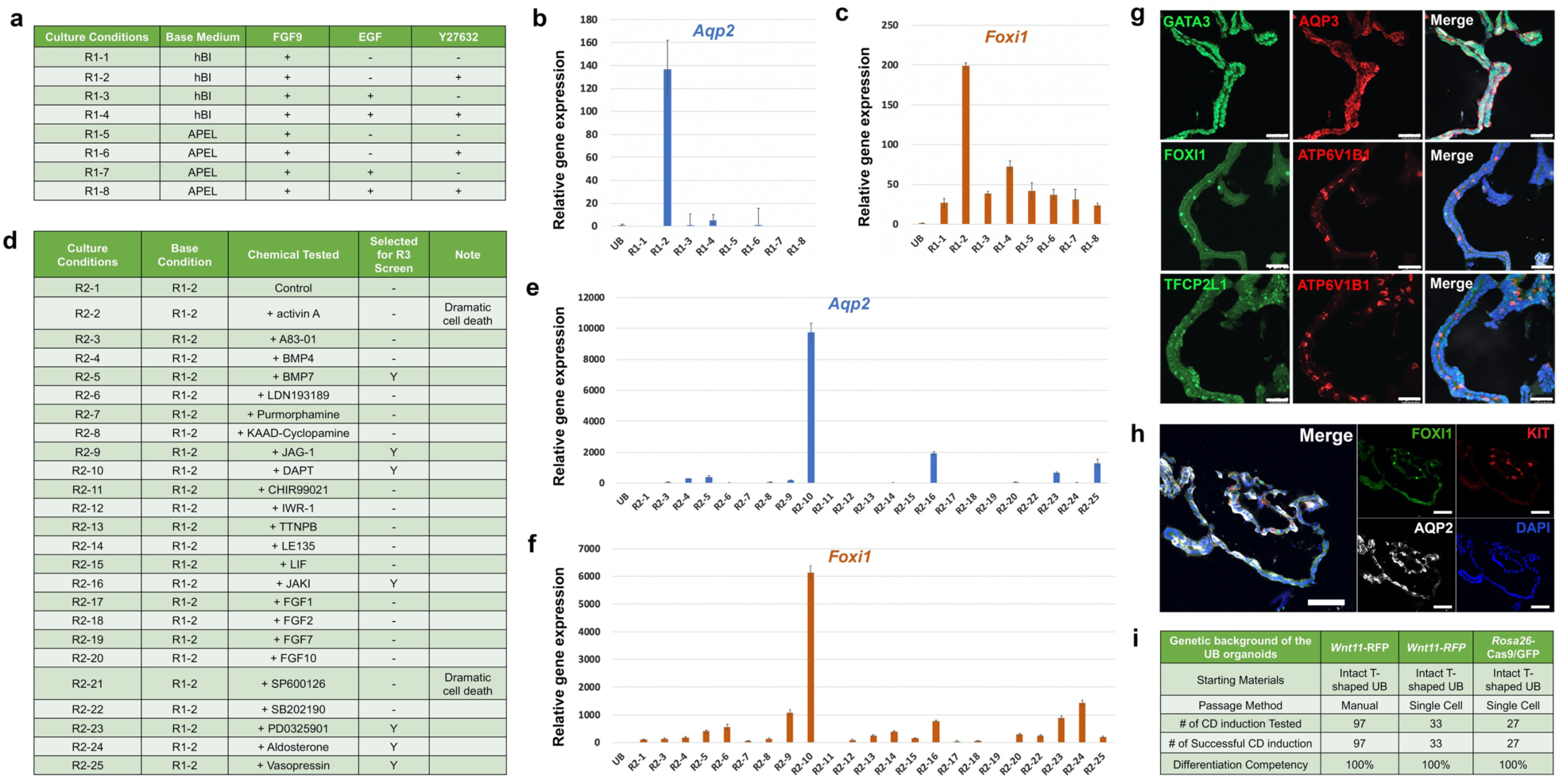
Expandable UB organoid-based screening for CD differentiation. **a**, Summary of conditions tested in the 1^st^ round of CD differentiation condition screening. **b, c**, qRT-PCR analyses of the 1^st^ round of CD differentiation condition screening for PC marker gene *Aqp2* (**b**, in blue) and IC marker gene *Foxi1* (**c**, in orange). Data are presented as mean ± s.d. Each column represents 3 technical replicates. **d**, Summary of chemicals tested in the 2^nd^ round of CD differentiation condition screening with the R1-2 medium as base medium. **e, f**, qRT-PCR analysis of the 2^nd^ round of CD differentiation condition screening for PC marker gene *Aqp2* (**e**, in blue) and IC marker gene *Foxi1* (**f**, in orange). Data are presented as mean ± s.d. Each column represents 3 technical replicates. Note that data for R2-2 and R2-21 were not presented here, because dramatic cell death were observed in those conditions and were thus excluded from this analysis. **g, h**, Immunostaining of cryo-section samples of differentiated CD organoids for ureteric lineage marker (GATA3) and various PC (AQP2 and AQP3) and IC markers (FOXI1, ATP6V1B1, and KIT), and TFCP2L1. Note the sporadic distribution of the IC in the organoid. Scale bar, 50 µm. **i**, Summary of CD organoid derivation efficiency.

**Extended Data Fig. 4.**
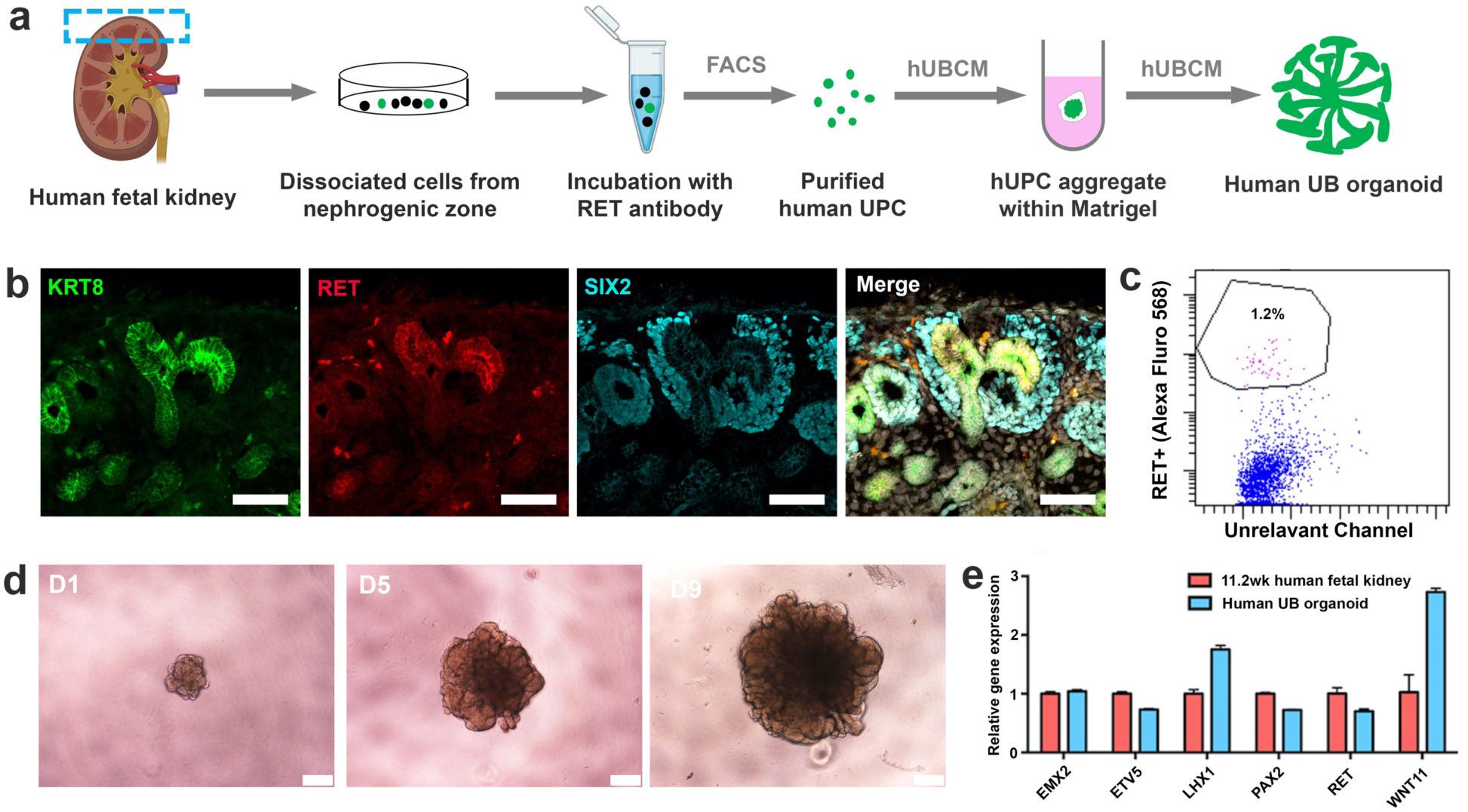
Generation of expandable human UB organoid from primary human UPCs. **a**, Schematic of the purification of primary human UPCs from the nephrogenic zone (illustrated as boxed region) of the human fetal kidney and the derivation of human UB organoid. **b**, Immunostaining of the human fetal kidney nephrogenic zone for UPC marker RET (red), broad UB lineage marker KRT8 (green), and NPC marker SIX2 (cyan). Scale bars, 50 µm. **c**, Representative flow cytometry analysis of RET+ UPC cells from dissociated human fetal kidney nephrogenic zone. **d**, Time course bright field images showing the growth of human UB organoid derived from primary human UPCs in a typical passage cycle. Scale bars, 200 µm. **e**, qRT-PCR analyses of human UB organoid derived from primary human UPCs for various UB markers. Human fetal kidney from 11.2-week (11.2wk) gestational age was used as control. Data are presented as mean ± s.d. Each column represents 3 technical replicates.

**Extended Data Fig. 5.**
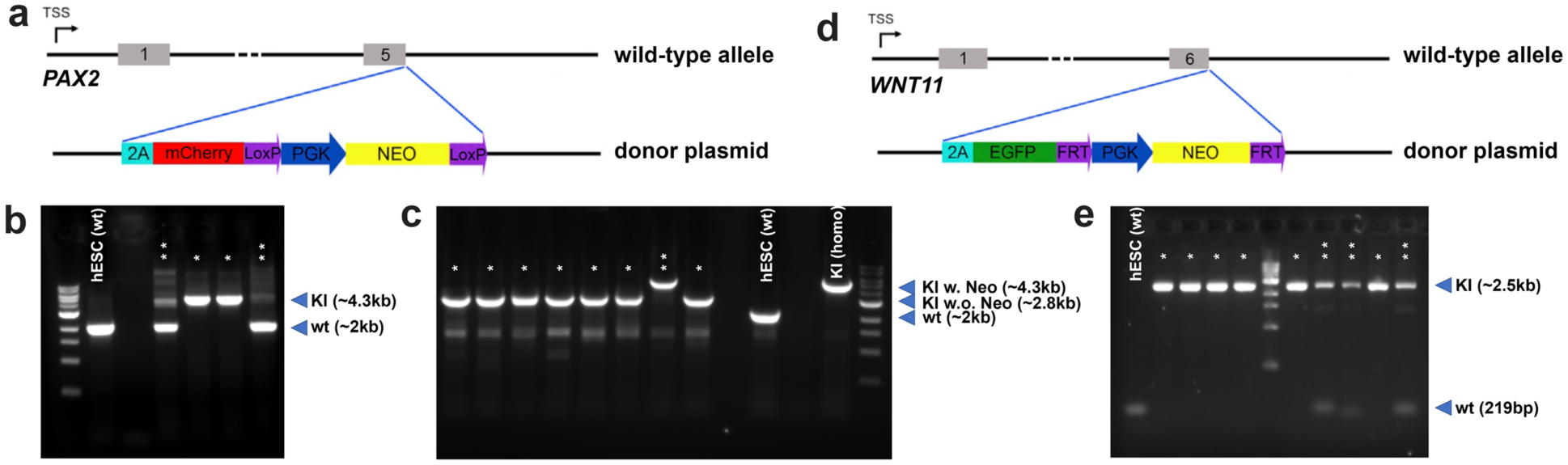
Genetic engineering hESC with a dual reporter system by CRISPR/Cas9. **a**, Schematic of the genetic engineering of *PAX2*-mCherry reporter into the hESC line. PGK-Neo cassette was used for antibiotic selection after gene targeting. This cassette was then excised from the genome by transient expression of Cre, before the second reporter system *WNT1*1-GFP was knocked in. **b**, PCR-based genotyping results of *PAX2*-mCherry reporter knockin. Wild-type (wt) PCR product is 2011bp, knockin (KI) product is 4308bp. *, biallelic KI clones; **, monoallelic KI clones; wild-type hESC “hESC (wt)”is used as a non-editing control. **c**, PCR-based genotyping result for Cre-based excision of PGK-Neo cassette. PCR product for *PAX2*-mCherry knockin with PGK-Neo excised (KI w.o. Neo) is 2779bp. *, clones with biallelic excision of PGK-Neo; **, clones without PGK-Neo excision; wild-type hESC “hESC (wt)”, and biallelic KI parental clone “KI (homo)” is used as controls. Note the very high efficiency in Cre-based PGK-Neo excision. **d**, Schematic of the engineering of *WNT11*-GFP reporter into the *PAX2*-mCherry reporter hESC line. PGK-Neo cassette was used for antibiotic selection after gene targeting. **e**, PCR-based genotyping result for *WNT11*-GFP reporter knockin. Wild-type (wt) PCR product is 219bp, knockin (KI) product is 2528bp. *, biallelic KI clones; **, monoallelic KI clones; wild-type hESC “hESC (wt)”is used as a non-editing control.

**Extended Data Fig. 6.**
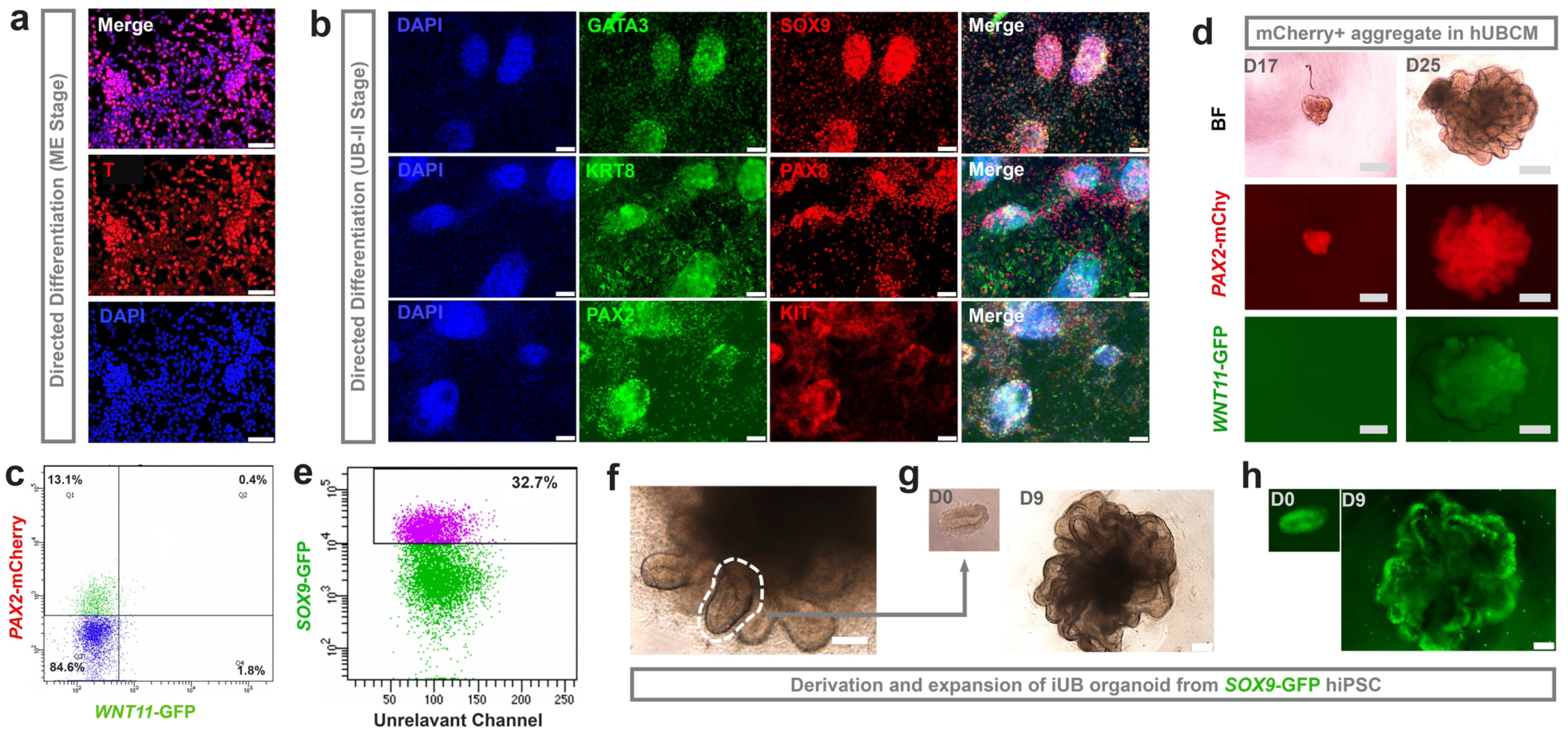
Generation of iUB organoids from various hPSC lines. **a**, Immunostaining of hESC-derived mesendoderm cells (at the end of ME stage) for mesendoderm marker Brachyury (T). Scale bars, 100 µm. **b**, Immunostaining of hESC-derived UB precursor cells (at the end of UB-II stage and prior to FACS sorting) for various UB lineage markers (KRT8, PAX8, PAX2, KIT, GATA3) and UB tip marker (SOX9). Scale bars, 100 µm. **c**, Flow cytometry analysis of mCherry+ and GFP+ cells differentiated from *WNT11*-GFP/*PAX2*-mCherry dual reporter hESCs. **d**, Bright field (BF) and fluorescence images showing the induction of *WNT11*-GFP expression in the mCherry+ aggregate upon extended culture in hUBCM. Scale bars, 200 µm. **e**, Flow cytometry analysis of GFP+ cells differentiated from *SOX9*-GFP reporter hiPSCs at the end of the UB-II stage. **f-h**, Bright field (**f,g**) and *SOX9*-GFP (**h**) images of branching iUB organoid derived from *SOX9*-GFP reporter hiPSCs in a typical passage cycle. The indicated budding structure shown in (**f**) was dissected and re-embedded into Matrigel for iUB expansion shown in (**g,h**). Scale bars, 200 µm.

**Extended Data Table 1.**
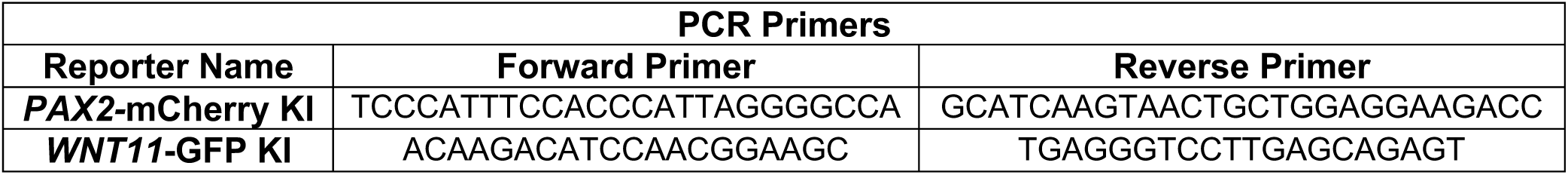
Genotyping primer sequences for detecting *PAX2*-mCherry and *WNT11*-GFP knockin (KI).

**Extended Data Table 2.**
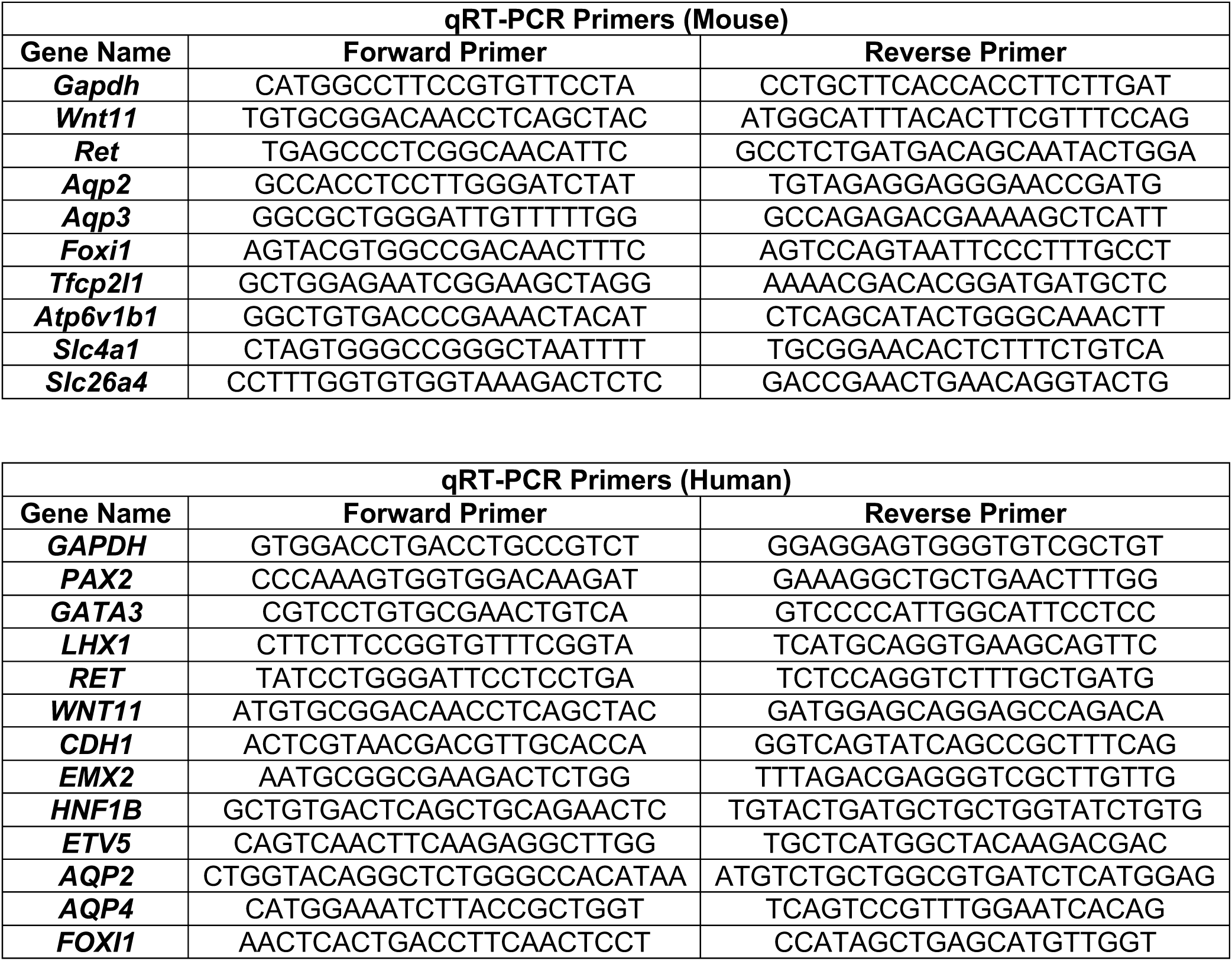
qRT-PCR Primer sequences.

**Extended Data Table 3.** Primary antibody information.

| **Antibodies** | **Company** | **Cat. No.** | **Dilution** |
| --- | --- | --- | --- |
| **KRT8** | DSHB | TROMA-I | 1:50 |
| **GATA3** | R&D Systems | AF2605 | 1:1000 |
| **SOX9** | Abcam | 185230 | 1:1000 |
| **RET (mouse)** | Cell Signaling Technology | 3223S | 1:300 |
| **RET (human)** | R&D Systems | AF1485 | 1:200 |
| **PAX2** | Covance | PRB-276P | 1:1000 |
| **PAX8** | Proteintech | 10336-1-AP | 1:1000 |
| **ETV5** | Abcam | Ab102010 | 1:300 |
| **CDH1** | BD | 610181 | 1:1000 |
| **AQP2** | Santa Cruz | sc-47710 | 1:300 |
| **AQP3** | Abcam | ab125219 | 1:300 |
| **FOXI1** | Novus Biologicals | NB300-926 | 1:300 |
| **ATP6V1B1** | Abcam | ab192612 | 1:300 |
| **TFCP2L1** | R&D Systems | AF5726 | 1:300 |
| **KIT** | Cell Signaling | 3074 | 1:300 |
| **SIX2** | Proteintech | 11562-1-AP | 1:1000 |
| **PODXL(mouse)** | R&D Systems | MAB1556 | 1:500 |
| **WT1** | Abcam | ab89901 | 1:1000 |
| **LTL** | Vector laboratories | B-1325 | 1:500 |
| **T** | R&D Systemss | AF2085 | 1:300 |

**Extended Data Table 4.** Growth factors and small molecules.

| **Reagent Name** | **Concentrations** | **Company** | **Cat. No.** |
| --- | --- | --- | --- |
| **LDN-193189** | 30nM, 200nM | Reagents Direct | 36-F52 |
| **TTNPB** | 0.1µM | TOCRIS | 0761 |
| **CHIR99021** | 3µM | Reagents Direct | 27-H76 |
| **JAK Inhibitor I** | 100nM | Stemcell Technologies | 74022 |
| **GDNF (mouse)** | 50ng/ml | PeproTech | 450-44-50µg |
| **GDNF (human)** | 50ng/ml | PeproTech | 450-10-50µg |
| **A83-01** | 0.2µM | STEMGENT | 04-0014 |
| **Rspo1** | 100ng/ml | R&D Systems | 4645-RS-100 |
| **SB202190** | 5µM | Axon Medchem | Axon 1304 |
| **Y27632** | 10µM | Cayman Chemical | 10005583 |
| **FGF9** | 50ng/ml | R&D Systems | 273-F9-025 |
| **FGF2** | 50,100, 200ng/ml | Stemcell Technologies | 78003 |
| **FGF7** | 50ng/ml | PeproTech | 100-19 |
| **FGF1** | 50ng/ml | Stemcell Technologies | 78187 |
| **FGF10** | 50ng/ml | PeproTech | 100-26 |
| **Aldosterone** | 100nM | Sigma-Aldrich | A9477 |
| **Vasopressin** | 1 I.U./ml | Sigma-Aldrich | V0377-100IU |
| **PD0325901** | 1µM | Reagents Direct | 39-C68 |
| **DAPT** | 5µM | Sigma-Aldrich | D5942 |
| **Activin A** | 10ng/ml, 50ng/ml | R&D Systems | 338-AC-010 |
| **EGF** | 50ng/ml | R&D Systems | 236-EG-200 |
| **BMP4** | 10ng/ml | R&D Systems | 314-BP-010 |
| **BMP7** | 10ng/ml | R&D Systems | 354-BP-010 |
| **Purmorphamine** | 1µM | STEMGENT | 04-0009 |
| **KAAD-Cyclopamin** | 50nM | STEMGENT | 04-0028 |
| **JAG-1** | 1µM | ANA SPEC | AS-61298 |
| **IWR-1** | 2.5µM | Sigma | 10161-5MG |
| **LE-135** | 0.5µM | Tocris | 2021 |
| **LIF** | 1000 units/ml | Millipore | ESG1107 |
| **SP600126** | 10µM | Tocris | 1496 |

#### Gene knockout

E11.5 mUB single cell suspension from *Rosa26*-Cas9/GFP background was used and lentiviral vectors were constructed using the lentiGuide-puro vector system (Addgene #52963) following standard protocol to make lentiviruses expressing three different gRNAs targeting GFP with the Cas9 cutting site 100-150bp apart, or three non-targeting gRNAs as control^5^. 100x concentrated lentivirus were used at 5X together with 10μM Polybrene diluted in mUBCM. 100µl virus-UBCM mixture was added to the U-bottom 96-well low-attachment plate well with 10 E11.5 mUBs that have been dissociated into single cells. The UB cells and virus were centrifuged together at 800g for 30 minutes for spin-infection. After the spin, virus-UBCM mixture was removed and fresh virus-UBCM mixture was added into the same well and the UB cells were spin-infected for another 30 minutes at 800g. Then, virus-UBCM mixture was removed and the infected UBs were washed three times with PBS, then aggregated and embedded in Matrigel and cultured in mUBCM in 37°C incubator following standard UB organoid culture procedures described above. 0.2μg/ml puromycin was applied to select for UB cells that have been successfully infected. The UB aggregate self-organize into typical branching organoid 4-5 days after infection.

## Culture Medium Recipes

### 1. mUBCM (mouse UB culture medium)

Basal medium: DMEM/F12 (1:1) (1X), Invitrogen, Cat. No. 11330-032.

Supplements:

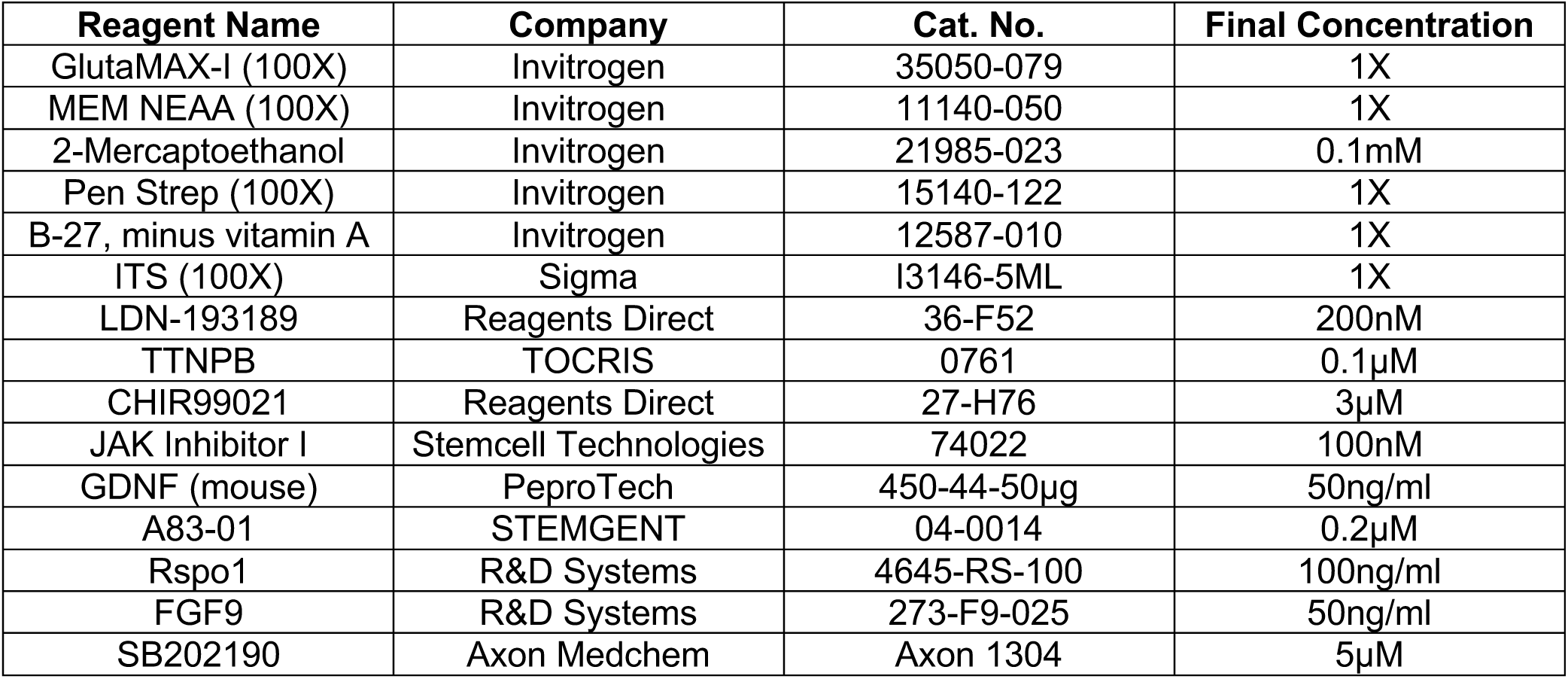

### 2. hUBCM (Human UB culture medium)

Basal medium: DMEM/F12 (1:1) (1X), Invitrogen, Cat. No. 11330-032.

Supplements:

| Reagent Name | Company | Cat. No. | Final Concentration |
| --- | --- | --- | --- |
| GlutaMAX-I (100X) | Invitrogen | 35050-079 | 1X |
| MEM NEAA (100X) | Invitrogen | 11140-050 | 1X |
| 2-Mercaptoethanol | Invitrogen | 21985-023 | 0.1mM |
| Pen Strep (100X) | Invitrogen | 15140-122 | 1X |
| B-27, minus vitamin A | Invitrogen | 12587-010 | 1X |
| ITS (100×) | Sigma | I3146-5ML | 1X |
| LDN-193189 | Reagents Direct | 36-F52 | 200nM |
| TTNPB | TOCRIS | 0761 | 0.1µM |
| CHIR99021 | Reagents Direct | 27-H76 | 3µM |
| JAK inhibitor I | Stemcell Technologies | 74022 | 100nM |
| GDNF (human) | PeproTech | 450-10-50µg | 50ng/ml |
| A83-01 | STEMGENT | 04-0014 | 0.2µM |
| Rspo1 | R&D Systems | 4645-RS-100 | 100ng/ml |
| FGF7 | PeproTech | 100-19 | 50ng/ml |
| SB202190 | Axon Medchem | Axon 1304 | 5µM |
| Y27632 | Cayman Chemical | 10005583 | 10µM |
| EGF | R&D Systems | 236-EG-200 | 50ng/ml |

### 3. CDDM (CD differentiation medium)

Basal medium: DMEM/F12 (1:1) (1X), Invitrogen, Cat. No. 11330-032.

Supplements:

| Reagent Name | Company | Cat. No. | Final Concentration |
| --- | --- | --- | --- |
| GlutaMAX-I (100X) | Invitrogen | 35050-079 | 1X |
| MEM NEAA (100X) | Invitrogen | 11140-050 | 1X |
| 2-Mercaptoethanol | Invitrogen | 21985-023 | 0.1mM |
| Pen Strep (100X) | Invitrogen | 15140-122 | 1X |
| B-27, minus vitamin A | Invitrogen | 12587-010 | 1X |
| ITS (100X) | Sigma | I3146-5ML | 1X |
| FGF9 | R&D Systems | 273-F9-025 | 50ng/ml |
| Y27632 | Cayman Chemical | 10005583 | 10µM |
| Aldosterone | Sigma-Aldrich | A9477 | 100nM |
| Vasopressin | Sigma-Aldrich | V0377-100IU | 1 I.U./ml |
| PD0325901 | Reagents Direct | 39-C68 | 1µM |
| DAPT | Sigma-Aldrich | D5942 | 5µM |

### 4. Stepwise directed differentiation to ND cells (D1 to D7 from hPSC)

Basal medium: DMEM/F12 (1:1) (1X), Invitrogen, Cat. No. 11330-032.

Supplements:

| Stages | Reagent Name | Final Concentration |
| --- | --- | --- |
| Common supplements for all three stages | GlutaMAX-I (100X) | 1X |
|  | MEM NEAA (100X) | 1X |
|  | 2-Mercaptoethanol (55mM) | 0.1mM |
|  | Pen Strep (100X) | 1X |
|  | B-27 Supplement (50X), minus vitamin A | 1X |
|  | ITS Liquid Media Supplement (100×) | 1X |
| ME Stage (D1-D3) | Activin A | 50ng/ml |
|  | CHIR99021 | 3μM |
| UB-I Stage (D3-D5) | FGF2 | 200ng/ml |
|  | TTNPB | 0.1μM |
|  | LDN-193189 | 30nM |
|  | A83-01 | 0.2μM |
| UB-II Stage (D5-D7) | FGF2 | 200ng/ml |
|  | TTNPB | 0.1μM |
|  | LDN-193189 | 30nM |

### 5. Mouse/Human kidney dissection medium

Basal medium: DMEM (1X), Invitrogen, Cat. No. 11995-040.

Supplements:

| Reagent Name | Company | Cat. No. | Final Concentration |
| --- | --- | --- | --- |
| GlutaMAX-I (100X) | Invitrogen | 35050-079 | 1X |
| MEM NEAA (100X) | Invitrogen | 11140-050 | 1X |
| 2-Mercaptoethanol | Invitrogen | 21985-023 | 0.1mM |
| Pen Strep (100X) | Invitrogen | 15140-122 | 1X |
| FBS | Gemini | 100-106 | 10% |

### 6. Mouse UB dissection medium

Basal medium: DMEM/F12 (1:1) (1X), Invitrogen, Cat. No. 11330-032.

Supplements:

| Reagent Name | Company | Cat. No. | Final Concentration |
| --- | --- | --- | --- |
| GlutaMAX-I (100X) | Invitrogen | 35050-079 | 1X |
| MEM NEAA (100X) | Invitrogen | 11140-050 | 1X |
| 2-Mercaptoethanol | Invitrogen | 21985-023 | 0.1mM |
| Pen Strep (100X) | Invitrogen | 15140-122 | 1X |
| B-27, minus vitamin A | Invitrogen | 12587-010 | 1X |
| ITS (100x) | Sigma | I3146-5ML | 1X |

